# Function of BriC Peptide in the Pneumococcal Competence and Virulence Portfolio

**DOI:** 10.1101/245902

**Authors:** Surya D. Aggarwal, Rory Eutsey, Jacob West-Roberts, Arnau Domenech, Wenjie Xu, Iman Tajer Abdullah, Aaron P. Mitchell, Jan-Willem Veening, Hasan Yesilkaya, N. Luisa Hiller

## Abstract

*Streptococcus pneumoniae* (pneumococcus) is an opportunistic pathogen that causes otitis media, sinusitis, pneumonia, meningitis and sepsis. The progression to this pathogenic lifestyle is preceded by asymptomatic colonization of the nasopharynx. This colonization is associated with biofilm formation; the competence pathway influences the structure and stability of biofilms. However, the molecules that link the competence pathway to biofilm formation are unknown. Here, we describe a new competence-induced gene, called *briC*, and demonstrate that its product promotes biofilm development and stimulates colonization in a murine model. We show that expression of *briC* is induced by the master regulator of competence, ComE. Whereas *briC* does not substantially influence early biofilm development on abiotic surfaces, it significantly impacts later stages of biofilm development. Specifically, *briC* expression leads to increases in biofilm biomass and thickness at 72h. Consistent with the role of biofilms in colonization, *briC* promotes nasopharyngeal colonization in the murine model. The function of BriC appears to be conserved across pneumococci, as comparative genomics reveal that *briC* is widespread across isolates. Surprisingly, many isolates, including strains from clinically important PMEN1 and PMEN14 lineages, which are widely associated with colonization, encode a long *briC* promoter. This long form captures an instance of genomic plasticity and functions as a competence-independent expression enhancer that may serve as a precocious point of entry into this otherwise competence-regulated pathway. Moreover, overexpression of *briC* by the long promoter fully rescues the *comE*-deletion induced biofilm defect *in vitro*, and partially *in vivo*. These findings indicate that BriC may bypass the influence of competence in biofilm development and that such a pathway may be active in a subset of pneumococcal lineages. In conclusion, BriC is a part of the complex molecular network that connects signaling of the competence pathway to biofilm development and colonization.

## Introduction

Bacteria form sessile communities termed biofilms, where they interact with each other to engage in collaborative and/or competitive behaviors (1). In *Streptococcus pneumoniae* (pneumococcus), these cell-cell interactions are commonly mediated by secreted peptides that interact with both producing and neighboring cells of the same species, and induce changes in gene regulation that result in altered phenotypes (2). These dynamic pneumococcal biofilms occur in chronic otitis media, chronic rhinosinusitis and nasopharyngeal colonization (3–8).

The ability to form biofilms is a critical component of pneumococcal disease (9). Biofilms serve as reservoirs for acute infections (10). In the middle ear, cells released from a biofilm are thought to be responsible for recurrent episodes of infection (4). Bacterial cells released from nasopharyngeal biofilms can seed pneumococcal transmission between individuals by being incorporated into nasal shedding. Alternatively, these cells can disseminate to tissues causing mild to severe diseases, such as otitis media, pneumonia, and sepsis (10). Pneumococcal cells released from biofilms display increased virulence relative to their planktonic or biofilm counterparts, suggesting that chronic biofilms set the stage for the stimulation of a virulence program activated upon the dispersal of cells (11). Moreover, pneumococci in a biofilm display decreased susceptibility to antibiotics, and are recalcitrant to treatment (6). Thus, biofilms are an important component of pneumococcal epidemiology in transmission, maintenance of asymptomatic colonization, and development of disease.

The transcriptional program required for the initiation and the growth of pneumococcal biofilms has been the subject of numerous investigations. It is clear that at least two quorum sensing (QS) signal transduction pathways are critical for biofilm development: competence and Lux (7,12–15). The competence pathway has been the subject of intense investigation for decades (16–28). Competence is activated by a classic two-component system where the extracellular <u>c</u>ompetence <u>s</u>timulating <u>p</u>eptide (CSP, encoded by *comC*) binds to the surface exposed ComD histidine kinase receptor, inducing its autophosphorylation and the subsequent transfer of the phosphate group to its cognate regulator, ComE (20,29). Activation of the competence pathway leads to increased expression of 5-10% of the pneumococcal genome in two main waves of gene expression (18,23). The first wave of induction is carried out directly by ComE; it upregulates a subset of competence genes (early genes) that include *comAB*, *comCDE*, as well as the alternative sigma factor, *comX*. The second wave of competence induction is regulated by ComX; it leads to an increase in the levels of at least 80 genes (late genes), that subsequently modulate important phenotypes such as transformation, metabolism, fratricide and biofilm formation (16,23,30). This competence program is upregulated during biofilm mode of growth *in vitro*, during interactions with human epithelial cells, and in lungs and brain after intranasal and intracranial challenges respectively in murine infection models (7,12,31). Importantly, in cell culture models, *comC* is required for biofilm development (12,14). Thus, activation of the competence pathway is important for productive biofilm formation and critical for pneumococcal infection and adaptation.

The Lux QS system also plays a role in biofilm formation. In this system, Lux QS is controlled by the AI-2 autoinducer, which is secreted and sensed by both Gram-positive and Gram-negative species. LuxS is a node in the regulation of competence, fratricide, and biofilm development (15,32). Lux upregulates competence via ComD and ComX (13). It contributes to bactericidal activity via upregulation of the choline binding murein hydrolase (CbpD). Through lysis, this bacteriocidal activity increases the levels of extracellular DNA, which is a key ingredient in the extracellular polymeric substance (EPS) that makes up the biofilm. Thus, the competence and Lux systems provide the molecular framework to coordinate multi-cellular bacterial communities to form and develop robust biofilms during infection.

Whereas the role of competence signaling in biofilm development is well established, the molecules that connect competence to biofilms are poorly understood (3,7,15,33). In this study, we identify one such molecule that links competence and biofilms. We characterize the gene encoding BriC (**<u>B</u>**iofilm **<u>r</u>**egulating peptide **<u>i</u>**nduced by **<u>C</u>**ompetence), a novel colonization factor in the competence pathway. Levels of *briC* are regulated by ComE, and *briC* promotes biofilm development and nasopharyngeal colonization. Further, genomic analysis of *briC* reveals polymorphisms in its promoter, where a subset of strains encode a RUP (for <u>r</u>epeat <u>u</u>nit of <u>p</u>neumococcus) sequence, which leads to additional and CSP-independent expression of *briC*.

## Results

### Identification of a competence-regulated Gly-Gly peptide

We have identified the gene encoding a putative secreted peptide that is co-regulated with competence (*spd_0391* (D39); *spr_0388* (R6); *sp_0429* (TIGR4)). Based on the results presented in this manuscript, we have termed it **<u>B</u>**iofilm-**<u>r</u>**egulating peptide **<u>i</u>**nduced by **<u>C</u>**ompetence (BriC). BriC was identified in our previously described *in silico* screen designed to capture cell-cell communication peptides in the pneumococcal genome (34). The known double glycine (Gly-Gly) streptococcal peptides are exported and proteolytically processed by dedicated ABC transporters that recognize N-terminal sequences with the Gly-Gly leader peptide (LSXXELXXIXGG)(20). In our previous work, we identified novel secreted pneumococcal peptides using a computational analysis to search for proteins with N-termini that contain a Gly-Gly leader. Our input set consisted of the alleles of two exported Gly-Gly peptides, the signaling molecule CSP and the bacteriocin BIP (20,35). Our output consisted of a position dependent probability matrix that captures the length and positional variability at each residue of the Gly-Gly motif. When we searched for this motif in a database of sixty streptococcal genomes, we defined a predicted secretome consisting of twenty-five sequence clusters, one of which corresponds to BriC (34).

To identify genes co-regulated with *briC*, we performed transcriptional studies using a NanoString probe set that reports on the abundance of the *briC* transcript as well as transcripts encoding a subset of pneumococcal regulators and cell wall proteins. We assessed the levels of *briC* transcript *in vitro* and *in vivo*. *In vitro* expression was measured by screening RNA extracted from mid-log planktonic cultures of a laboratory strain (R6-derivative (R6D)). *In vivo* expression was evaluated by analysis of middle-ear effusions recovered from chinchillas infected with a clinical PMEN1 strain. The mRNA levels of the *briC* were positively associated with *comC* and *comE in vitro* (strain R6D: R^2^=0.61 and 0.79, respectively) *and in vivo* (strain PN4595-T23: R^2^=0.92 and 0.88, respectively). It is noteworthy that changes in the expression of genes in this locus were observed in the studies that first documented the competence regulon (18,23). Specifically, Peterson and colleagues observed changes in *briC* levels, however the association between *briC* and CSP was below the statistical threshold (23). Further, Dagkessamanskaia and colleagues observed an upregulation in the gene downstream of *briC*, predicted to be in the same operon (18). Given that in strains R6, R6D, and D39, this downstream gene is truncated, this study does not explore the function of the downstream gene. In summary, our gene expression analysis suggests that *briC* is induced by competence.

To directly test whether *briC* is a competence-regulated peptide, we employed a fusion of the *briC* promoter to the *lacZ* reporter (R6 P*briC-lacZ*). Stimulation of the signal transduction system that initiates competence by addition of CSP1 led to an induction of the β-galactosidase activity by about twenty-five-fold (**Fig. 1**). Induction of the *briC* promoter was specific to the CSP pherotype encoded by strain R6. The β-galactosidase activity was observed upon addition of CSP1, the CSP pherotype from strain R6, but not upon addition of the non-cognate CSP2 pherotype (**Fig. 1**). Thus, we conclude that *briC* is a competence-responsive gene.

**Fig. 1.**
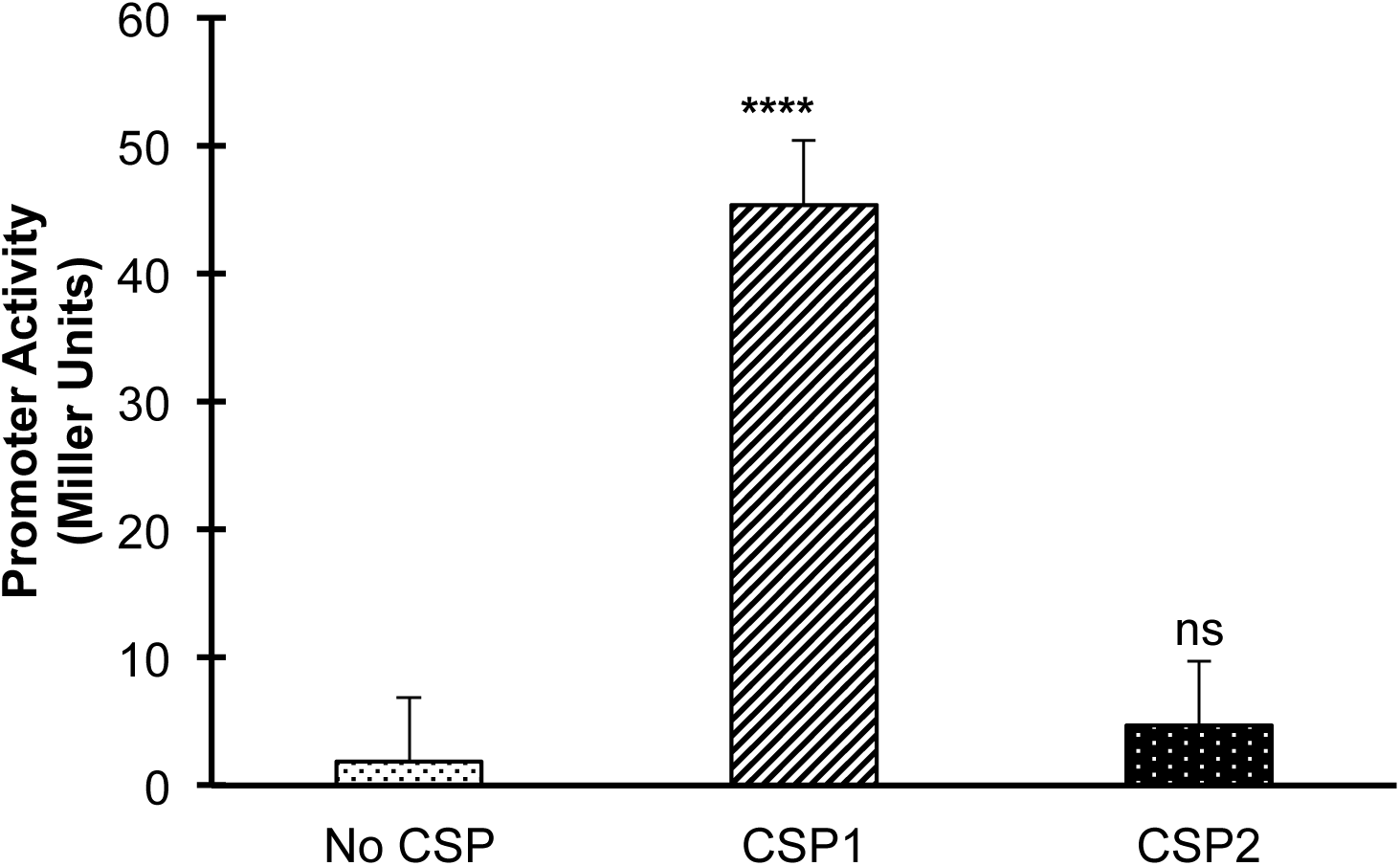
Expression of *briC* is induced by cognate CSP. β-galactosidase assay measuring P*briC-*lacZ activity in pneumococcal R6 cells grown to exponential phase in Columbia Broth at pH 6.6 followed by no treatment or treatment with CSP1 or CSP2 for 30 minutes. Y-axis denotes P*briC*-*lacZ* expression levels in Miller Units. Activity is expressed in nmol p-nitrophenol/min/ml. Error bars represent standard error of the mean for biological replicates *(at least n=3)*; **** *p*<0.0001 using ANOVA followed by Tukey’s post-test.

### Levels of *briC* transcripts are directly regulated by ComE

Our *in silico* analysis of the *briC* promoter in strains R6 and R6D revealed the presence of a ComE-binding site. ComE binds a well-defined sequence consisting of two imperfect direct repeats of nine nucleotides separated by a gap of twelve or thirteen base pairs (36). Our analysis of the putative *briC* promoter across pneumococcal strains revealed an excellent match to the ComE-binding box (**Fig. 2A**). To further investigate the association between ComE and *briC*, we tested whether CSP-induction of *briC* requires ComE. We compared the CSP-induction of *briC* in a wild-type (R6D WT) strain to that of an isogenic *comE*-deletion mutant (R6DΔ*comE*), using qRT-PCR analysis. In WT cells, the addition of CSP triggered a significant increase in levels of *briC* at 10 minutes post-addition, with levels slowly decreasing by 15 minutes (**Fig. 2B**). This trend follows the temporal pattern observed for the levels of *comE* that has been associated with genes under direct controls of ComE (18,23). In contrast, the transcript levels of *briC* were unaffected by CSP addition in the Δ*comE* strain, indicating that the expression of *briC* requires ComE (**Fig. 2B**). These results strongly suggest that *briC* is directly regulated by ComE. In addition, ComE plays a critical role in controlling transformation, thus we investigated whether *briC* impacts transformation efficiency (**Supplementary Results and Fig. S1**). We find that absence of *briC* leads to only a minor decrease in transformation efficiency.

**Fig. 2.**
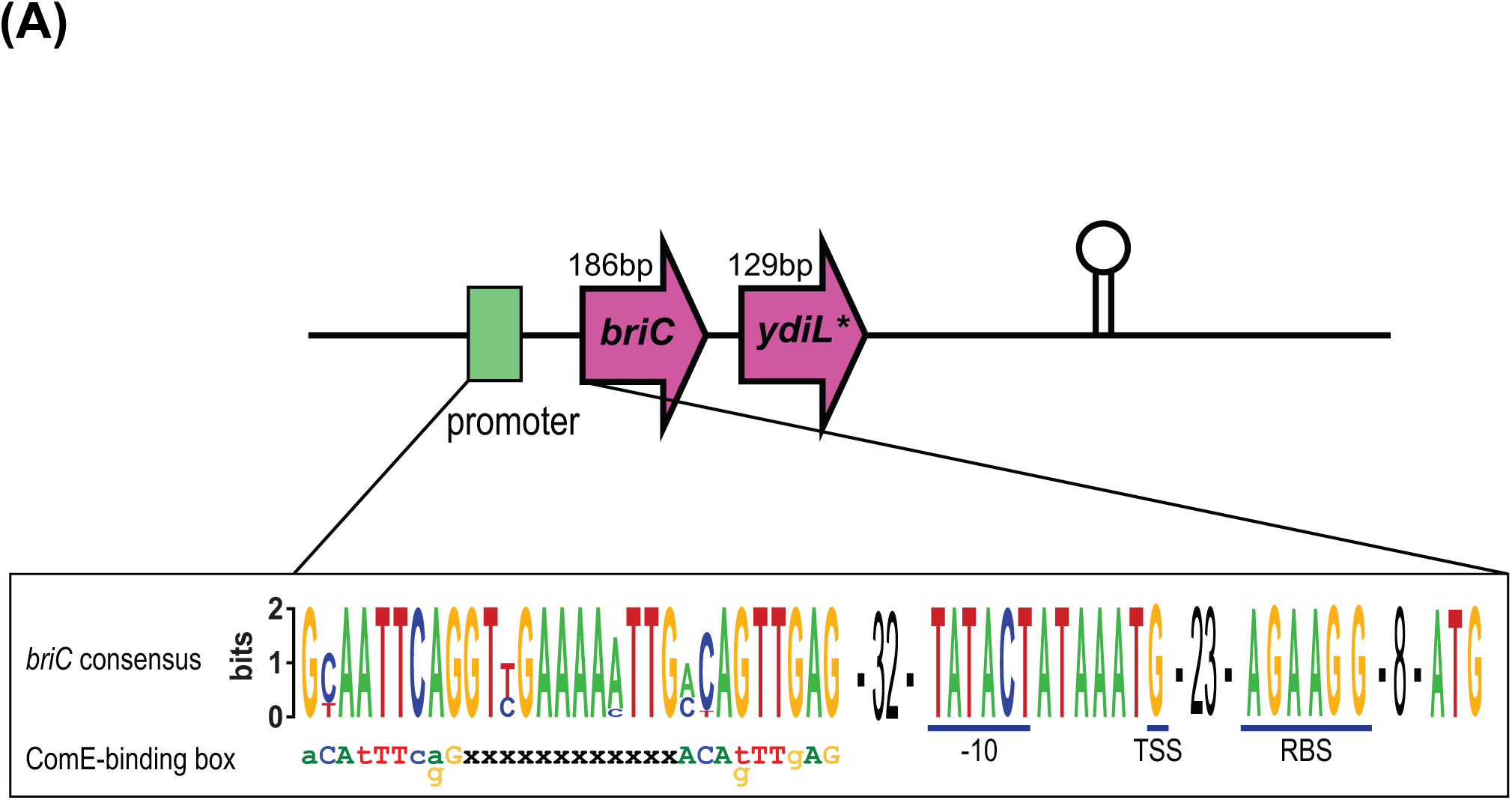
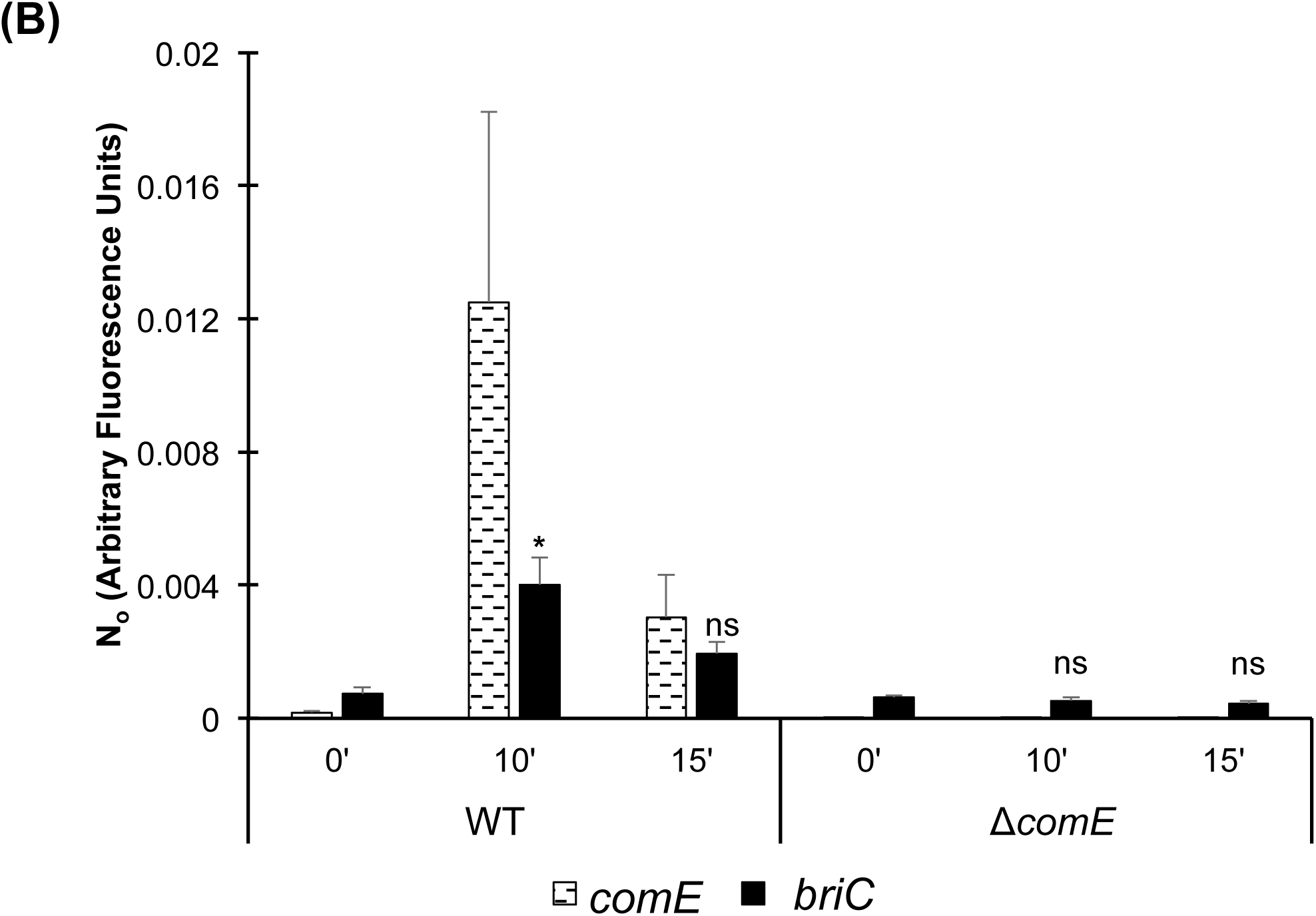
CSP-induction of *briC* is ComE-dependent. (A) Genomic organization of the *briC* locus, displaying a ComE-binding box. Green: ComE-binding box within the *briC* promoter region. The expanded region denotes a logo of ComE-binding box generated from thirty-four pneumococcal genomes represented in Figure 4A. This consensus is aligned with the published ComE-binding box consensus sequence (Ween et al., 1999). The putative −10 region, the **t**ranscription **s**tart **s**ite (TSS) as determined by Cappable-Seq (Slager et al., 2018), the **r**ibosome **b**inding **s**ite (RBS) and the transcriptional terminator are labeled. The downstream gene is predicted to be a pseudogene in R6D, R6 and D39. In TIGR4, this region encodes two coding sequences (SP_0430 and SP_0431). The R6D sequence corresponds to the C-terminal of SP_0430. (B) mRNA transcript levels of *briC* (solid black) and *comE* (dashed black lines) as measured by qRT-PCR in R6D WT & R6DΔ*comE* cells. Cells were grown in Columbia broth at pH 6.6 to an OD_600_ of 0.3, and then treated with CSP1 for either 0’, 10’ or 15’. Data was normalized to 16S rRNA levels. Y-axis denotes normalized concentrations of mRNA levels in arbitrary fluorescence units as calculated from LinRegPCR. Error bars represent standard error of the mean calculated for biological replicates *(n=3)*; ‘ns’ denotes non-significant, * *p*<0.05 using ANOVA followed by Tukey’s post-test relative to the respective 0’ CSP treatment. Further, *briC* levels are also significantly higher in WT relative to Δ*comE* cells for the same time points post-CSP treatment (*p*<0.05).

### BriC plays a key role in biofilm development

To investigate whether expression of *briC* plays a role in biofilm development, we compared biofilm formation across WT (R6D WT), *briC* deletion mutant (R6DΔ*briC*), and *briC* complemented (R6DΔ*briC*::*briC*) strains grown on an abiotic surface at 24h and 72h post-seeding. No difference was observed in biofilm biomass and thickness at 24h post-seeding, suggesting that expression of *briC* does not contribute to early stages of biofilm formation (**Fig. 3A, B**). In contrast, at 72h post-seeding, Δ*briC* biofilms displayed significantly reduced biomass and thickness when compared to WT (**Fig. 3C, D**). Further, biofilms with Δ*briC::briC* cells restored the WT phenotype at this time-point (**Fig. 3C, D**). The indistinguishable biofilm parameters of WT and ∆*briC* cells at 24h post-seeding suggests that there is no fitness-related growth difference between the strains and indicates that the biofilm defect is biologically relevant. These findings suggest that *briC* contributes to late biofilm development.

**Fig. 3.**
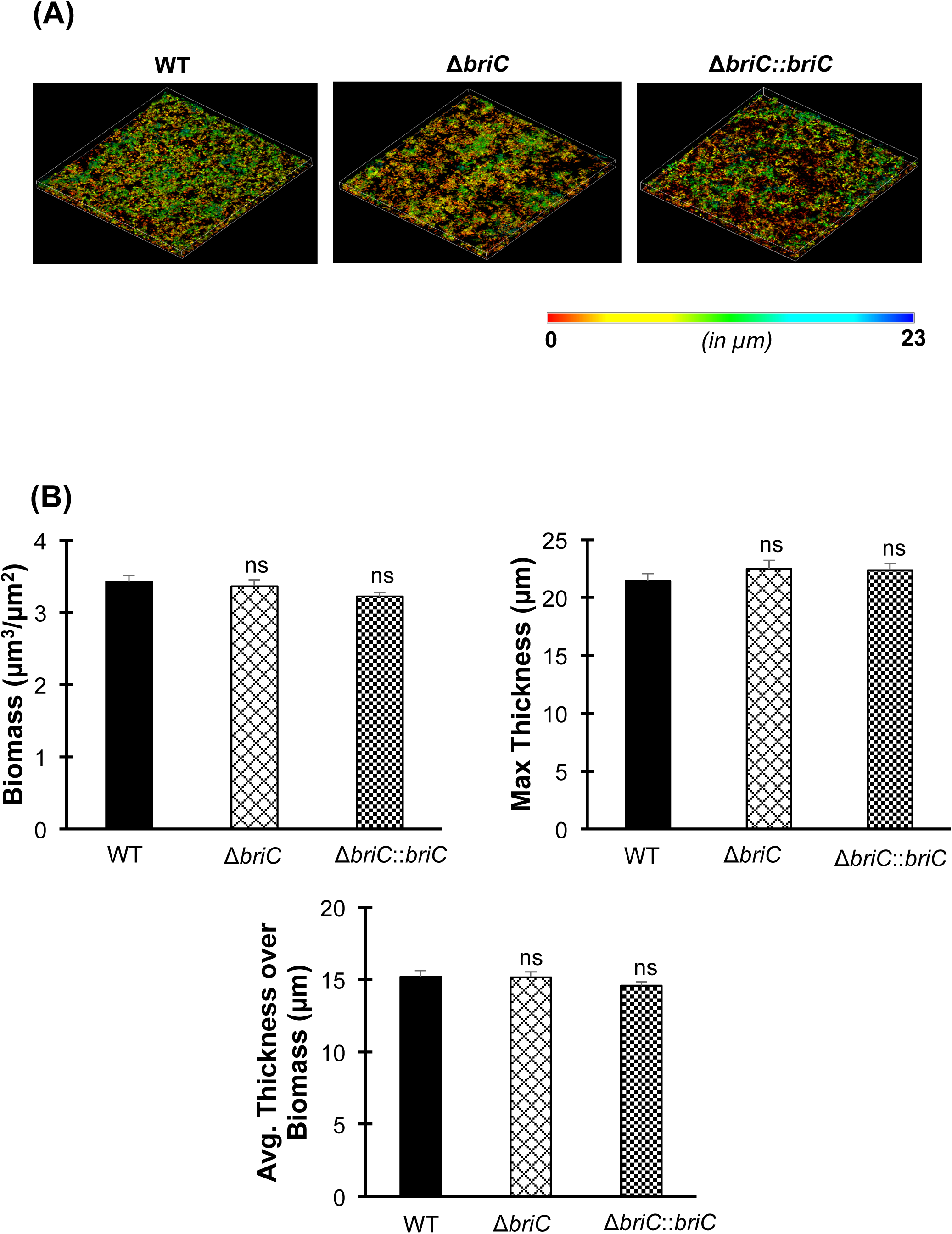
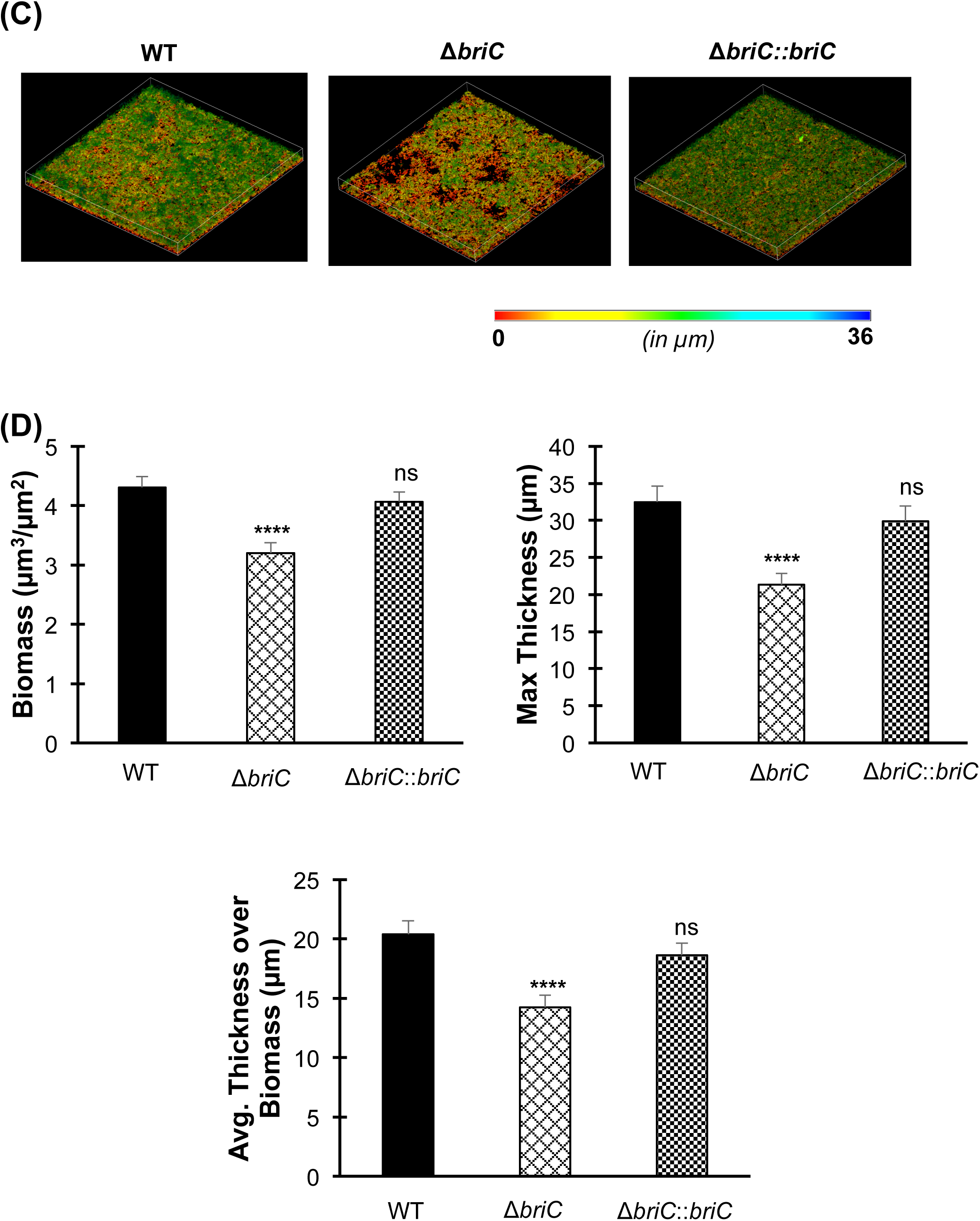
BriC stimulates late biofilm development. Representative confocal microscopy images showing top view of the reconstructed biofilm stacks of WT, Δ*briC* and Δ*briC::briC* cells of strain R6D stained with SYTO59 dye at **(A)** 24-hr, and **(C)** 72-hr. Images are pseudo-colored according to depth (scales shown). COMSTAT2 quantification of **(B)** 24-hr, and **(D)** 72-hr biofilm images. Y-axis denotes units of measurement: μm^3^/μm^2^ for biomass, and μm for maximum thickness and average thickness over biomass. Error bars represent standard error of the mean calculated for biological replicates *(n=3)*; “ns” denotes non-significant comparisons, **** *p*<0.0001 using ANOVA followed by Tukey’s post-test.

### *briC* is widely distributed across pneumococcal strains

To investigate the prevalence of *briC*, we investigated its distribution across the genomes of pneumococcus and related streptococci. To place the distribution in the context of phylogeny, we used a published species tree generated from a set of fifty-five genomes (34,38) (**Table S1**). The genomes encompass thirty-five pneumococcal genomes that span twenty-nine multi-locus sequence types as well as eighteen serotypes and nontypeable strains; eighteen genomes from related streptococcal species that also colonize the human upper respiratory tract, namely *S. pseudopneumoniae*, *S. mitis*, and *S. oralis*; and finally, two distantly related *S. infantis* strains as an outgroup. In this set, all the pneumococcal genomes encode *briC*, and there are two highly similar alleles (labeled allele 1A and 1B, **Fig 4A, B**). Further, we identified four additional alleles in the related streptococci (**Fig. 4A, File S1**). Next, we extended this analysis to a set of 4,034 pneumococcal genomes available in the pubMLST database (these correspond to all the genomes with at least 2Mb of sequence) (39). In total, 98.5% (3,976 out of 4,034) of these genomes encode a *briC* allele, suggesting *briC* is highly prevalent across pneumococcal strains. We find that alleles 1A and 1B are prominent in this larger set, with 1,824 and 1,187 representatives respectively. After manual curation, we retrieved nineteen distinct *briC* alleles across these pneumococcal genomes (**Fig. 4B**). Six of the polymorphic residues are located in the putative leader sequence. The conserved region of the leader sequences corresponds to the amino acids preceding the Gly-Gly (20,34), thus the polymorphisms at the N-terminal end of the BriC sequence are not predicted to influence export. One polymorphic residue replaces the glycine in the Gly-Gly motif with a serine; it seems probable that this variation may influence processing and/or export. In addition, position −2 from the C-terminus encodes either an alanine or a threonine. This variation is at the C-terminus, predicted to be the functional end of the molecule, such that the difference in size and polarity at this residue is likely to impact structure or binding of BriC to its targets. The *briC* gene is induced by competence, so we investigated whether there is a correlation between CSP pherotypes and alleles encoding BriC. However, we did not find these to be associated (**Table S2**). Finally, to investigate the distribution of *briC* in other species and genera, we used BLASTp to search the non-redundant database (40). We find that BriC homologs are present in strains of related streptococci, *S. pseudopneumoniae*, *S. mitis* and *S. oralis*, but we did not identify homologues in more distant species. The phylogenetic distribution of *briC* supports a conserved role across pneumococci and a subset of related streptococcal species.

**Fig. 4.**
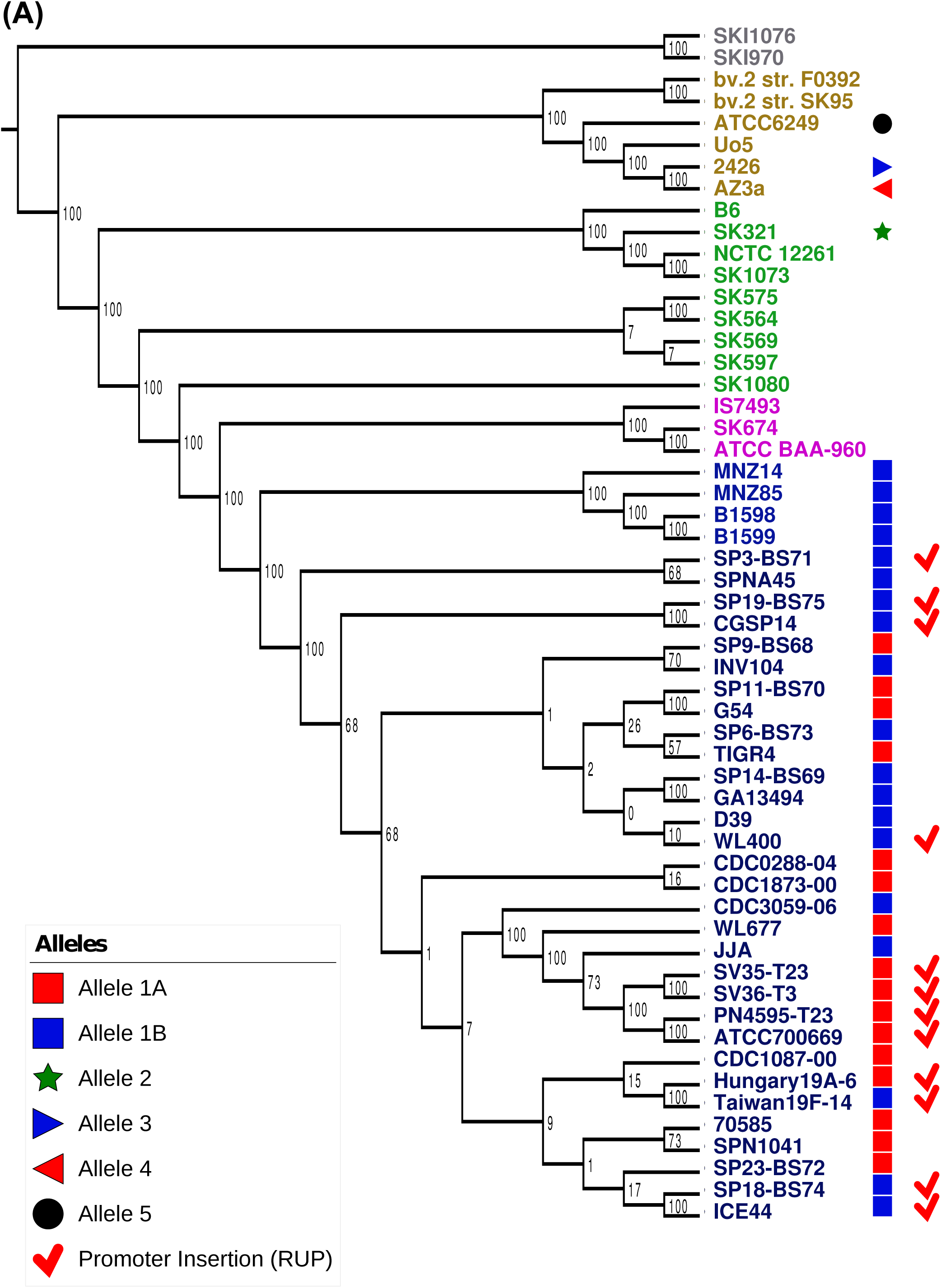
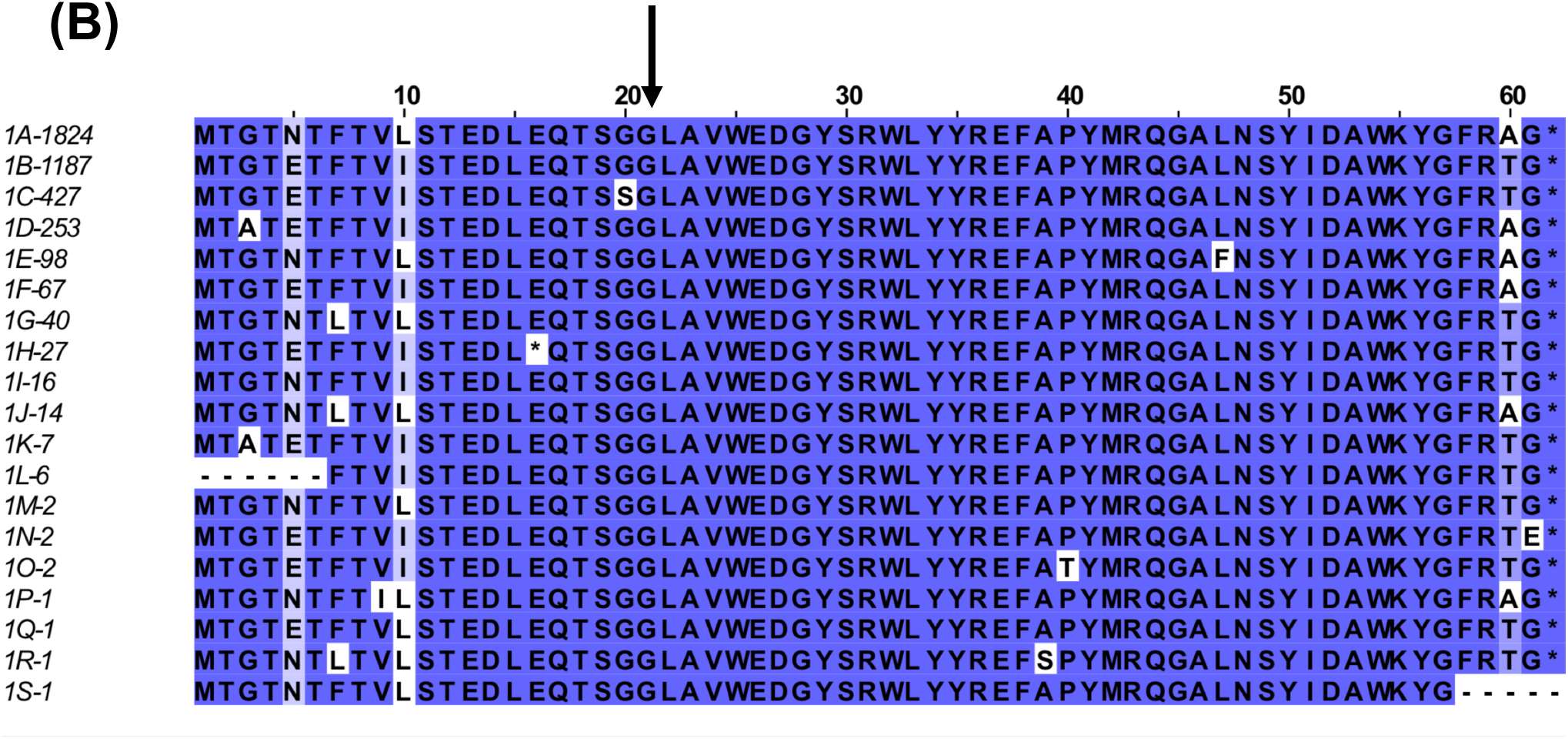
Distribution of the genomic region encoding BriC across streptococcal strains. **(A)** Distribution of *briC* alleles in fifty-five streptococcal genomes. The *briC* alleles are visualized against a maximum likelihood tree of streptococcal genomes generated from the core genome, where the numbers on the branches represent bootstrap values. Different species in the tree are color-coded as follows: *S. pneumoniae* (blue), *S. pseudopneumoniae* (pink), *S. mitis* (green), *S. oralis* (beige), and *S. infantis* (grey). The shapes at the tip of the branches illustrate *briC* alleles. Types 1A and 1B represent variants of the alleles widespread across pneumococcal strains; types 3–5 denotes alleles outside the species. The red tick denotes strains that have a long *briC* promoter due to a RUP insertion. In PMEN1 strains, this variant leads to increase in basal levels of *briC* in a CSP-independent manner. **(B)** Alignment of 19 BriC alleles identified in the database of 4,034 pneumococcal genomes. Alleles are labeled 1A-1S followed by the number of representatives in the database (total 3,976). Sequences are colored based on percent identity to highlight the variability between alleles. Black arrow denotes the predicted cleavage site.

### Inter-strain differences in the *briC* promoter are associated with diverse regulation of *briC* in clinically important lineages

Our analysis of the promoter region of *briC* in our curated set of genomes reveals that a subset of strains encode for a 107 bp insertion within the region upstream of *briC* (**Fig. 4**). The additional nucleotides are located after the ComE-binding site and before the transcriptional start site, and correspond to a **r**epeat **u**nit in **p**neumococcus (RUP) sequence (41,42). RUP is an insertion sequence derivative with two clear variants, which may still be mobile (42). The RUP sequence upstream of *briC* corresponds to RUPB1.

In our curated genomes, the long RUP-encoding promoter is present in multiple strains, including those from the clinically important PMEN1 and PMEN14 lineages (**Fig. 4A**). Using our expanded database of 4,034, we determined that the vast majority of the PMEN1 and PMEN14 genomes encode a long promoter. The high prevalence of the long promoter in these lineages suggests that this form was present in the ancestral strains from these lineages and/or provides a fitness advantage in these genomic backgrounds.

To investigate how this genomic difference influences *briC* expression, we generated a LacZ reporter strain. The 263bp upstream of *briC* from the PMEN1 strain, PN4595-T23, were fused to *lacZ* to produce the P*briC*_long_-*lacZ* reporter, and its reporter activity was compared to that of the P*briC*-*lacZ* generated with the fusion of 159bp upstream of *briC* obtained from strain R6. To control for the possibility that the influence of the RUP sequence might be strain-dependent, we tested these reporter constructs in the R6 and the PMEN1 backgrounds, in both the absence and presence of CSP treatment (**Fig. 5A, B)**. The presence of RUP dramatically increased the basal levels of *briC* in the absence of CSP, and this increase was observed in both R6 and PMEN1. Furthermore, both constructs were induced upon the addition of CSP. These findings suggest that the RUP sequence serves as an expression enhancer; it increases the levels of *briC* transcripts and this increase is CSP-independent. Thus, in some lineages, *briC* appears to be under the control of both CSP-dependent and CSP-independent regulation.

**Fig. 5.**
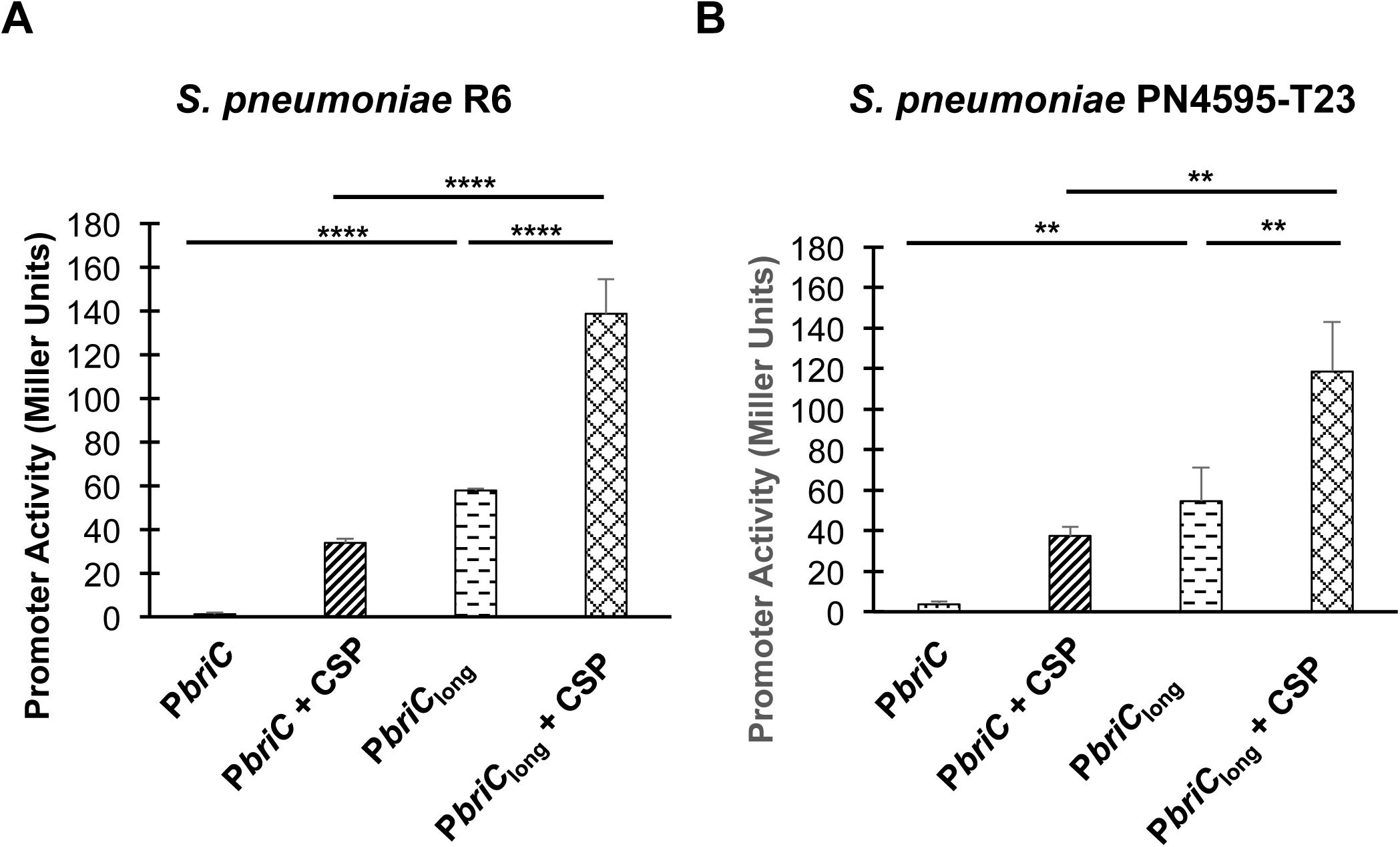
Long *briC* promoter is associated with an increase in the basal levels of *briC*. β-galactosidase assay comparing the LacZ activity of the R6 (short promoter, P*briC-lacZ)* and PN4595-T23 (long promoter with RUP, P*briC*_long_*-lacZ)* promoters. Both promoter activities were tested in **(A)** strain R6 and **(B)** strain PN4595-T23. Cells were grown in Columbia broth at pH 6.6 until mid-log phase, followed by either no treatment or treatment with CSP for 30 minutes. Y-axis denotes promoter activity in Miller Units expressed in nmol p-nitrophenol/min/ml. Error bars represent standard error of the mean for biological replicates *(n=3)*; ** *p*<0.01, & **** *p*<0.0001 using ANOVA followed by Tukey’s post-test.

### Expression of *briC* driven by the long promoter bypasses the impact of competence induction in biofilm development

Next, we investigated the biological impact of the natural variations in the *briC* promoter on biofilm development. It has been well established that competence promotes biofilm development. Specifically, deletion of the *comC* (encodes CSP) and *comD* (encodes histidine kinase of competence TCS) genes lead to a reduction in *in vitro* biofilms in strains D39 and TIGR4 (7,12). In this study, we have established that *briC* also promotes biofilm development (**Fig. 3B**), and that the RUP-containing long promoter serves as an expression enhancer (**Fig. 5**). Thus, we hypothesized that expression of *briC* from the long promoter may bypass the impact of competence in biofilm development.

First and in concurrence with previous work, we observed that a strain with a *comE* deletion (R6DΔ*comE*) displays a reduction in biofilm biomass and thickness relative to the WT strain (**Fig. 6A, B**). ComE influences the expression of numerous genes. To determine whether the biofilm defects were primarily due to its impact on *briC* induction, we tested a construct with a disruption of the ComE-binding box in the *briC* promoter (Δ*briC::*P*briC*_Shuffled ComE-box_-*briC*). This strain displays a significant reduction in biofilm biomass and thickness relative to the WT strain (**Fig. 6A, B).** Moreover, no difference was observed in the biofilm parameters for both of these mutants, suggesting that the absence of *briC* expression is a contributor to the *in vitro* biofilm defect in the *comE* deletion mutant. Next, we determined that a strain with increased basal levels of *briC* driven by the RUP-containing long promoter (R6DΔ*comE*::P*briC*_long_-*briC)* fully rescued the biofilm defect observed in R6DΔ*comE* (**Fig. 6A, B**). In addition, increased expression of *briC* in the wild type background (R6D WT::P*briC*_long_-*briC)* did not lead to a significant increase in biofilm biomass and thickness relative to the wild type (**Fig. 6A, B**). Together, these data strongly suggest that *briC* is a critical molecular link between competence and biofilm formation, and that natural variations in the *briC* promoter are physiologically relevant.

**Fig. 6.**
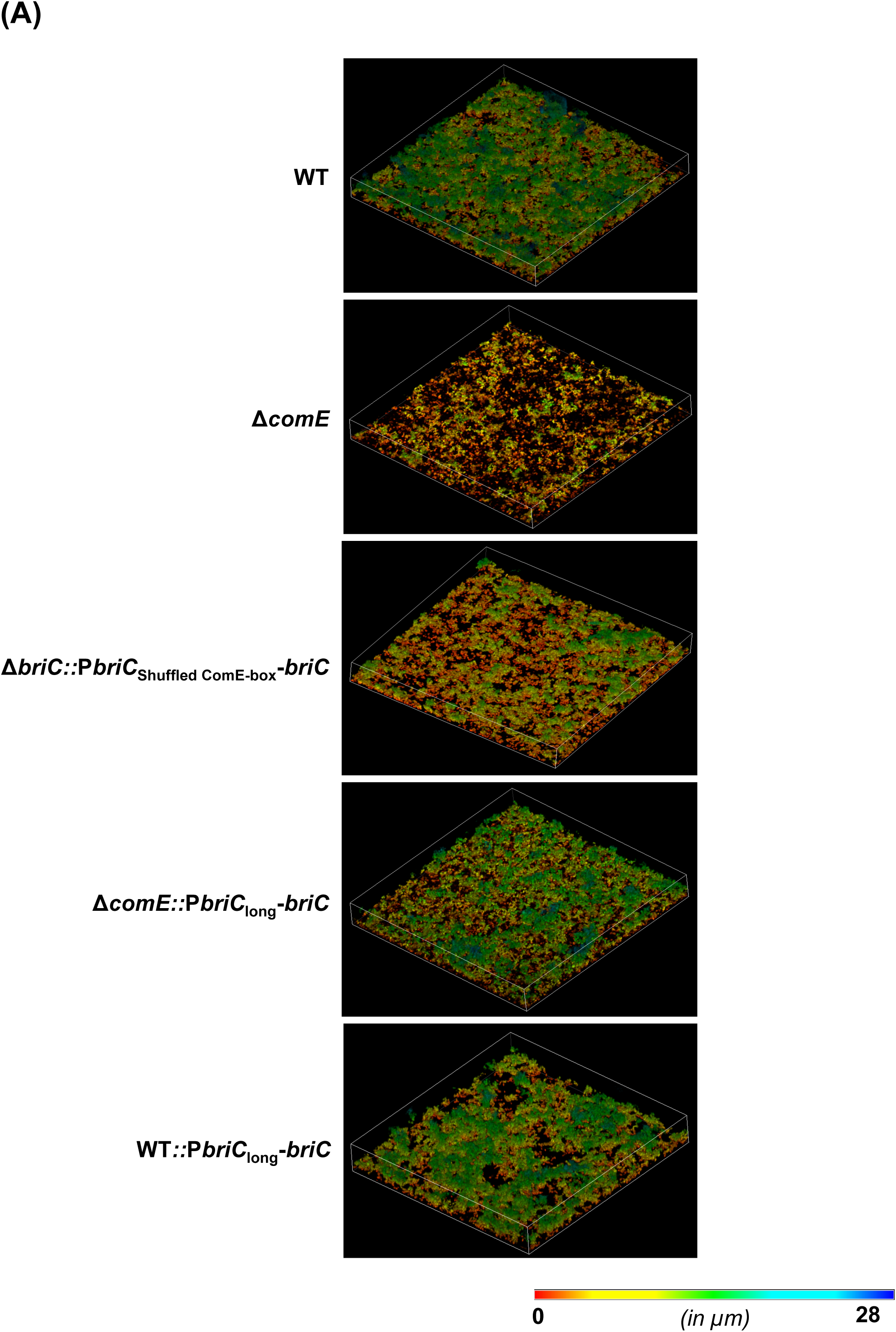
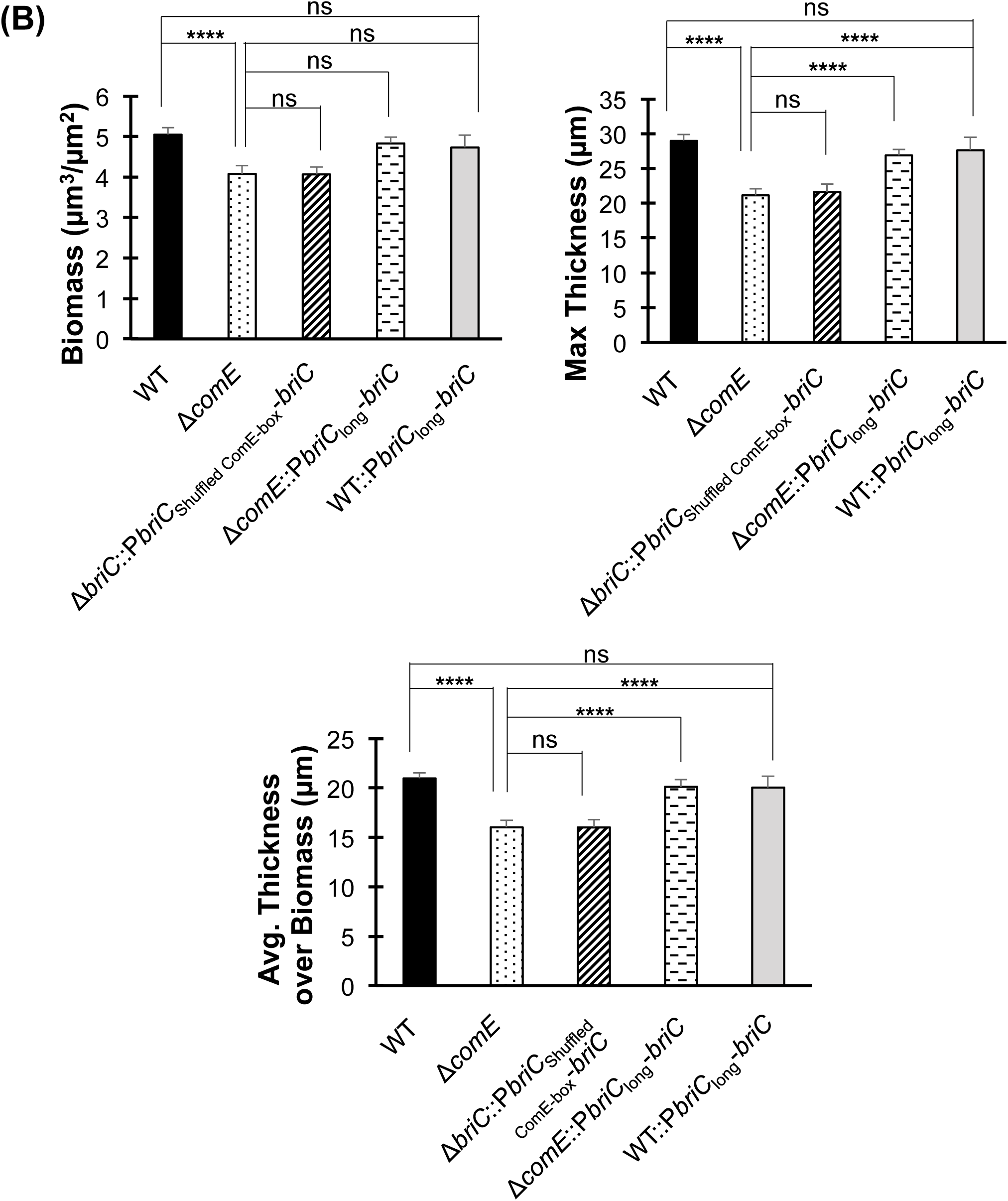
BriC plays a pivotal role in regulating biofilm development. **(A)** Representative confocal microscopy images showing top view of the reconstructed biofilm stacks of WT, Δ*comE,* Δ*briC::*P*briC*_Shuffled ComE-box_-*briC*, Δ*comE::*P*briC*_long_-*briC* and WT::P*briC*_long_-*briC* cells of strain R6D stained with SYTO59 dye at 72-hr. Images are pseudo-colored according to depth (scale shown). **(B)** COMSTAT2 quantification of 72-hr biofilm images. Y-axis denotes units of measurement: μm^3^/μm^2^ for biomass, and μm for maximum thickness and average thickness over biomass. Error bars represent standard error of the mean calculated for biological replicates *(at least n=3)*; “ns” denotes nonsignificant comparisons, and **** *p*<0.0001 using ANOVA followed by Tukey’s post-test.

### ComAB plays a role in secretion of the protein encoded by *briC*

Since BriC is associated with the competence pathway and is able to rescue the biofilm defects associated with competence signaling, we investigated whether competence associated transporters play a role in exporting BriC. In pneumococcus, the ComAB and BlpAB C39-peptidase transporters export peptides with a Gly-Gly leader (43–45). These transporters recognize the N-terminal leader of target sequences, and cleave these sequences at the Gly-Gly motif (45,46). In strains R6 and R6D, BlpAB is not functional due to a frameshift mutation that leads to an early stop codon (47). Thus, we hypothesized that as a Gly-Gly peptide co-expressed with genes of the competence pathway, BriC may be exported via the ComAB transporter. We tested this hypothesis in two ways. First, we measured whether deletion of *comAB* influences the ability of a strain with competence-independent expression of *briC* to rescue the Δ*comE*-biofilm defect. Second, we compared secretion of a BriC reporter construct in a WT strain with that in a *comAB* deletion mutant.

Our biofilm data suggests that ComAB plays a role in transporting BriC. At 72h post-seeding, a *comE/comAB*-double deletion mutant strain expressing *briC* from the RUP-encoding long promoter (R6DΔ*comE*Δ*comAB::*P*briC*_long_-*briC*) displayed a biofilm with reduced biomass and thickness when compared to the equivalent construct in a *comE*-deletion background (R6DΔ*comE::*P*briC*_long_-*briC*) (**Fig. 7A, B**). Hovewer, the biofilm levels were not reduced to the levels observed in the Δ*comE* strain. These results suggest that under these conditions, ComAB may not be the only transporter that contributes to the export of BriC.

**Fig. 7.**
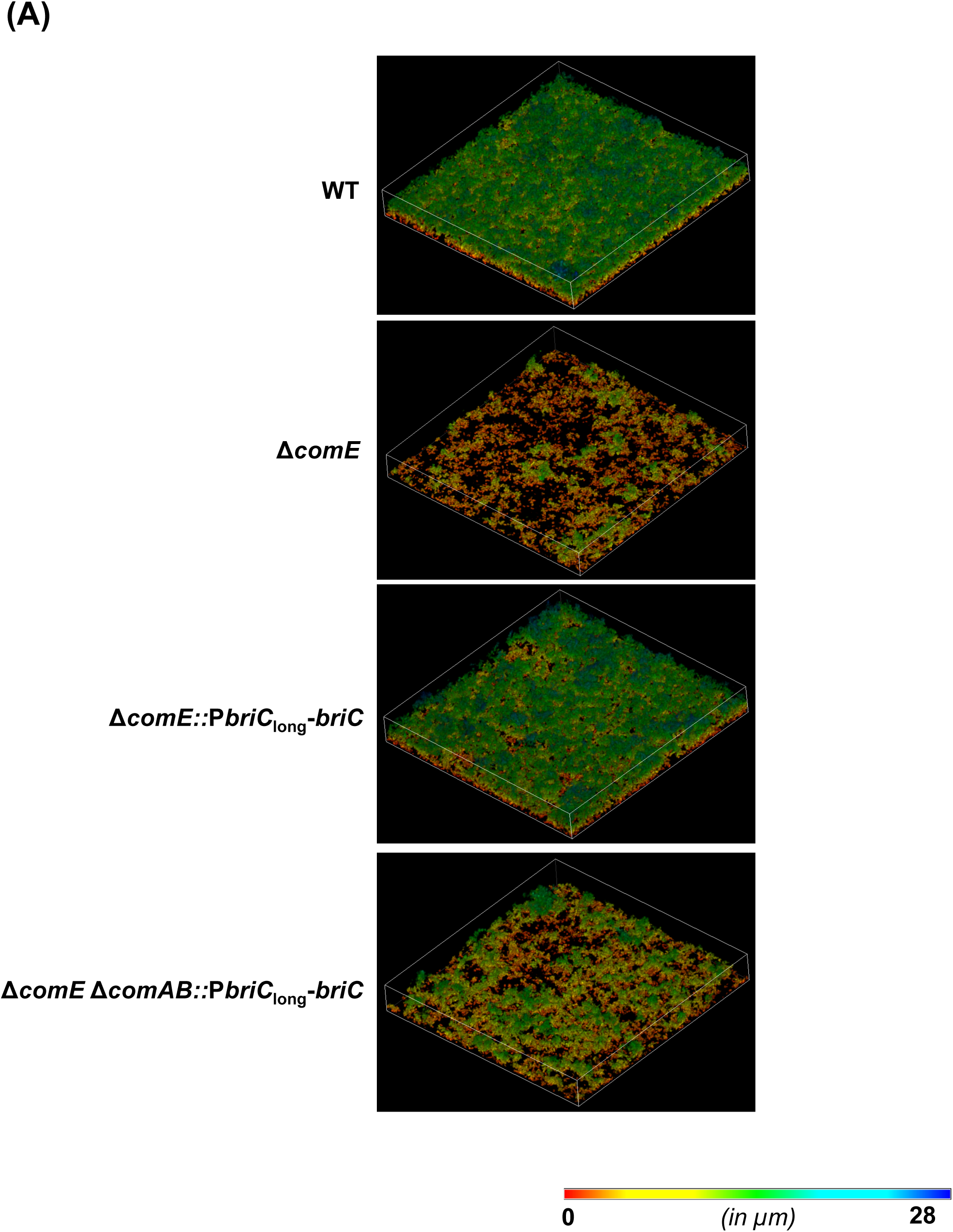
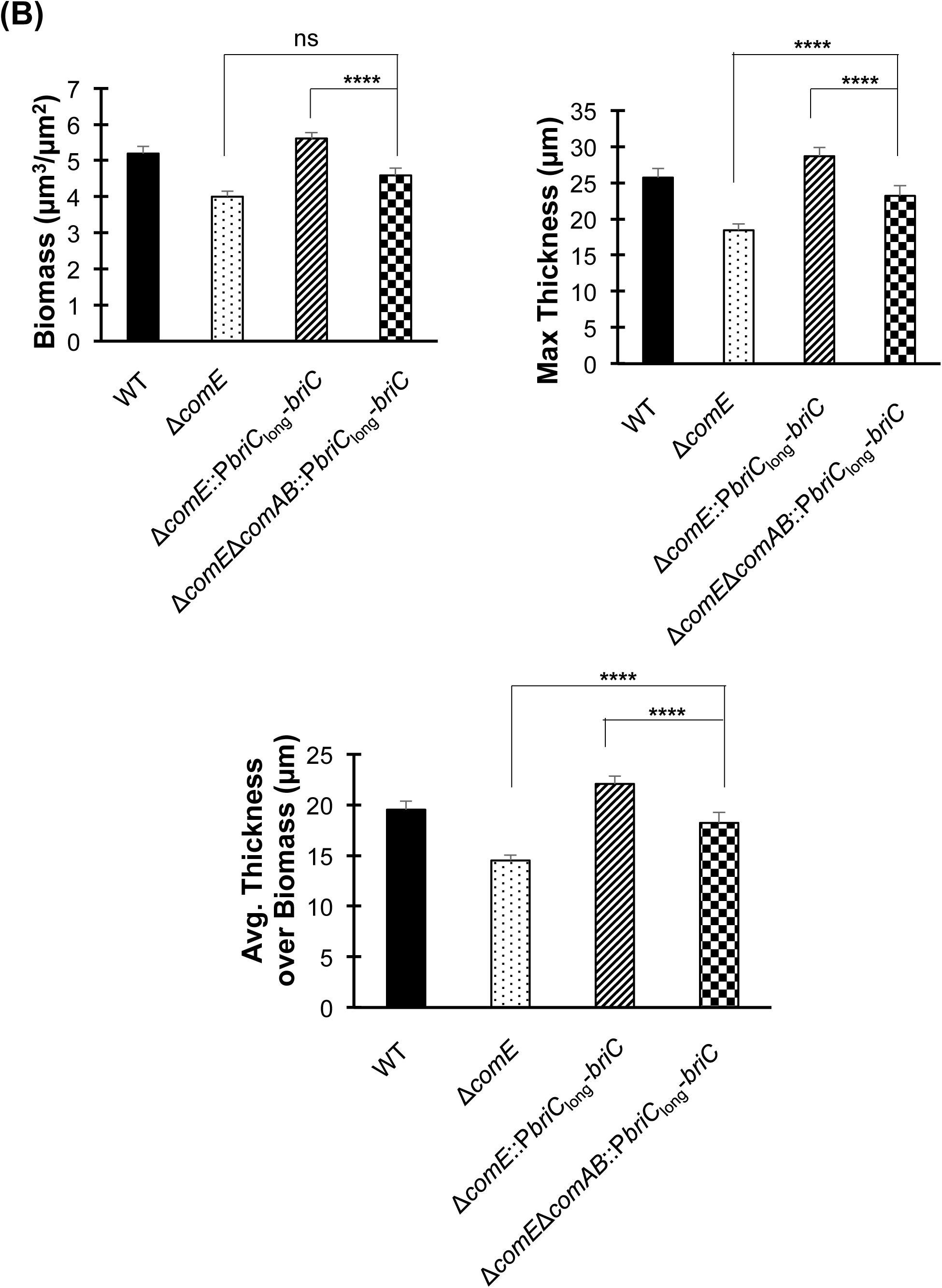
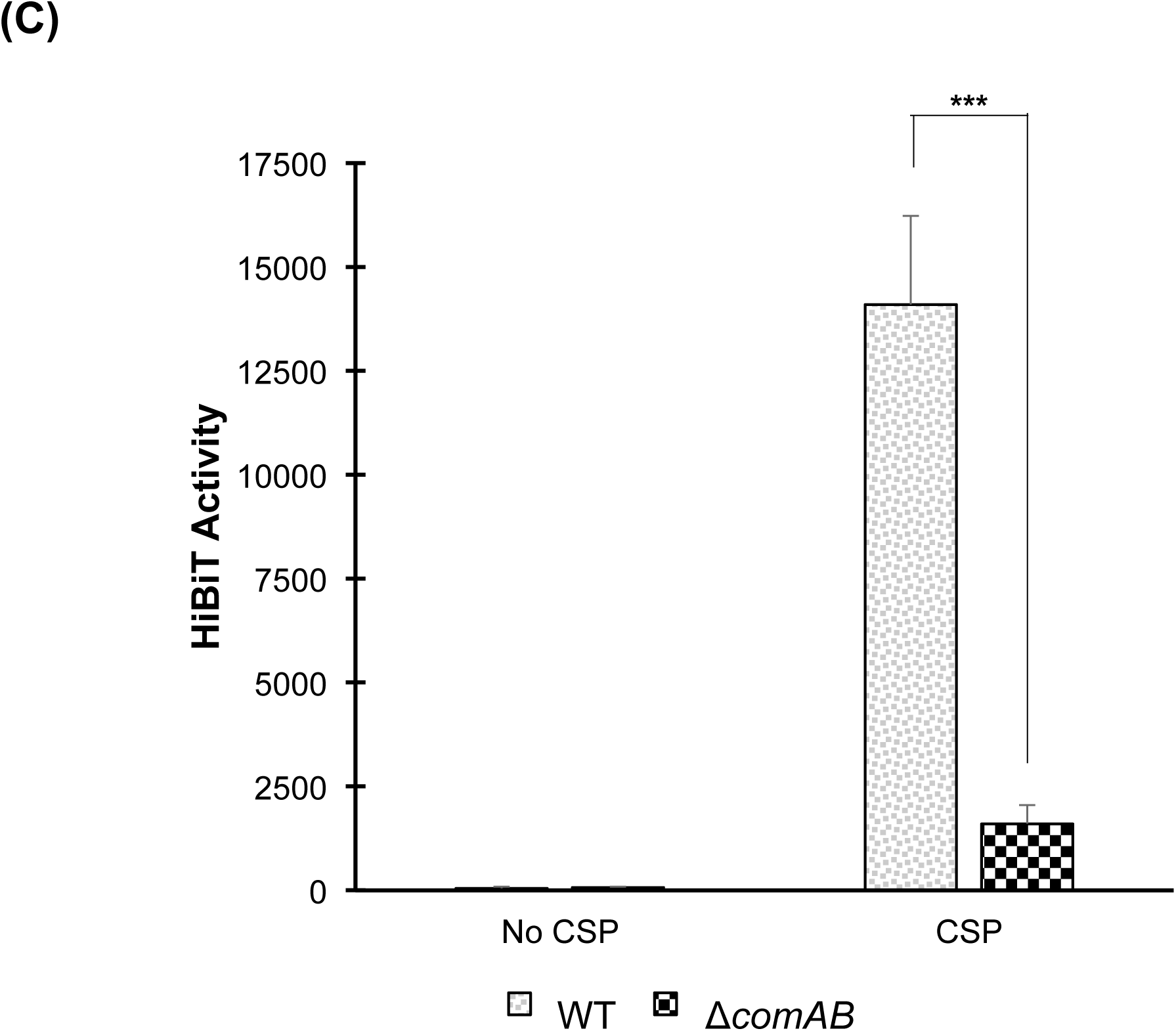
ComAB plays a role in the export of BriC. **(A)** Representative confocal microscopy images showing top view of the reconstructed biofilm stacks of WT, Δ*comE,* Δ*comE::*P*briC*_long_-*briC* and Δ*comE*Δ*comAB::*P*briC*_long_-*briC* cells of strain R6D stained with SYTO59 dye at 72-hr. Images are pseudo-colored according to depth (scale shown). **(B)** COMSTAT2 quantification of 72-hr biofilm images. Y-axis denotes units of measurement: μm^3^/μm^2^ for biomass, and μm for maximum thickness and average thickness over biomass. Error bars represent standard error of the mean calculated for biological replicates *(atleast n=3).* **(C)** Extracellular Nano-Glo HiBiT activity of the BriC reporter produced by WT and Δ*comAB* cells (whole cells). The HiBiT activity was measured by recording luminescence with an integration time of 2000 milliseconds. Error bars represent standard deviation calculated for biological replicates *(n=3)*; “ns” denotes non-significant comparisons, *** *p*<0.001, and **** *p*<0.0001 using ANOVA followed by Tukey’s post-test.

To further elucidate the role of ComAB in the export of BriC, we employed the HiBiT tag detection system, which was recently used to detect secretion of BlpI (44). The HiBiT tag corresponds to an 11-residue peptide. The assay works by addition of an inactive form of luciferase (LgBit) to the extracellular milieu. When both LgBit and HiBiT combine, they generate bioluminescence (48,49). To study BriC transport, we fused the putative BriC leader sequence to the HiBiT tag and expressed this reporter under the control of the native (short) *briC* promoter in WT and Δ*comAB* R6-strains. We measured the extracellular bioluminescence produced by this reporter both in the presence and absence of CSP (**Fig. 7C, Table S3**). In the absence of CSP, the levels of secreted HiBiT resembled that of background (WT cells without HiBiT), consistent with very low expression of N-terminal BriC-HiBiT as well as low expression of the ComAB transporter. In the WT background, upon addition of CSP, N-terminal BriC-HiBiT is induced and the extracellular level of HiBiT is significantly increased, consistent with HiBiT export. In contrast, in the Δ*comAB* background, upon addition of CSP, N-terminal BriC-HiBiT is induced but the extracellular levels of HiBiT do not substantially increase, consistent with lack of HiBiT export. Combined, these results strongly suggest that ComAB serves as a transporter for BriC.

### BriC is important for *in vivo* colonization

During nasopharyngeal colonization, pneumococci form biofilms and upregulate the competence pathway. Thus, we investigated the role of *briC* in nasopharyngeal colonization using an experimental murine colonization model. Our *in vitro* investigations have been performed using strain R6D strain, which is defective in colonization due to the absence of a capsule. Thus, we performed colonization experiments with the serotype 2 strain, D39, which is the ancestor of strain R6 (50). Mice were colonized with D39 WT, the *briC*-deletion mutant (D39Δ*briC*) or the *briC*-complemented (D39Δ*briC*::*briC*) strains. Comparison of the number of bacteria in nasal lavages immediately after inoculation revealed that mice in the three cohorts received the same number of bacteria. In contrast, nasal lavages at three and seven days post-inoculation revealed decreased levels of D39Δ*briC* relative to WT in the nasal wash (**Fig. 8A**). Furthermore, the WT levels were restored in the complemented strain (**Fig. 8A**). These findings indicate that *briC* encodes a novel colonization factor.

**Fig. 8.**
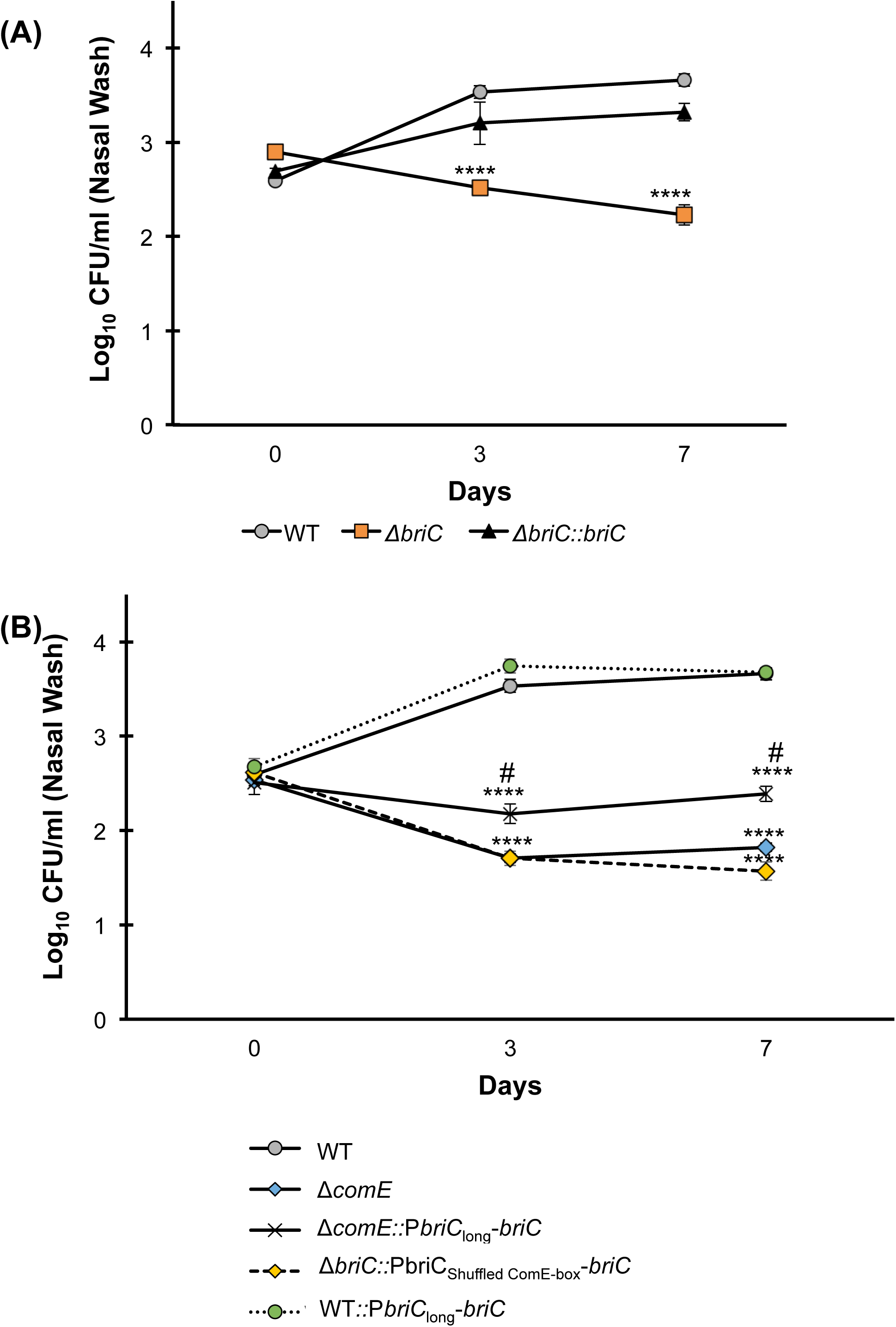
BriC contributes to pneumococcal colonization of the mouse nasopharynx. CD1 mice were infected intranasally with 20μl PBS containing approximately 1 × 10^5^ CFU of **(A)** WT (grey circles), Δ*briC* (orange squares), and Δ*briC::briC* (black triangles) **(B)** WT (grey circles), Δ*comE* (blue diamonds), and Δ*comE::PbriC*_long_-*briC* (black crosses), Δ*briC::*P*briC*_Shuffled ComE-box_-*briC* (yellow triangles), and WT::*PbriC*_long_-*briC* (green circles) cells of the pneumococcal strain D39. At predetermined time points (0, 3 & 7 days post-infection), at least five mice were culled, and the pneumococcal counts in the nasopharyngeal washes were enumerated by plating on blood agar. Y-axis represents Log_10_ counts of CFU recovered from nasal washes. X-axis represents days post-inoculation. Each data point represents the mean of data from at least five mice. Error bars show the standard error of the mean. **** *p*<0.0001 relative to the WT strain, and # *p*<0.0001 relative to the Δ*comE* strain, calculated using ANOVA and Tukey post-test.

In *in vitro* biofilms, *briC* links competence to biofilms. First, disruption of the ComE-binding box in the *briC* promoter led to a biofilm defect similar to that observed in a Δ*comE* strain. Second, overexpression of *briC* driven by the long version of the promoter was found to restore the competence-dependent defect in *in vitro* biofilm development. Thus, we investigated the behavior of these strains in the pneumococcus colonization model. We found that the strain with a disruption of the ComE-binding box (Δ*briC::*P*briC*_Shuffled ComE-box_-*briC*) within the *briC* promoter was defective for colonization, the decreased bacterial counts resembled those in the Δ*comE* strain (**Fig. 8B**). These findings suggest that *briC* is a substantial mediator of the role of ComE on colonization. Further, addition of this long *briC* promoter to Δ*comE* cells (D39Δ*comE::*P*briC*_long_-*briC)* partially rescues the colonization defect of the D39Δ*comE* strain. That is, the numbers of bacterial cells of strain D39Δ*comE::*P*briC*_long_-*briC* recovered from the nasal lavages at both three and seven days post-inoculation were significantly higher than the numbers of D39Δ*comE* cells recovered, but less than that of the D39 WT (**Fig. 8B)**. Finally, the overexpression of *briC* in the WT background (D39 WT::P*briC*_long_-*briC*) does not impact colonization. Thus, we conclude that BriC is a contributor to the competence-induced stimulation of nasopharyngeal colonization observed in strain D39. Further, natural variations leading to a long *briC* promoter appear to dampen the impact of competence in colonization.

## Discussion

An important component of pneumococcal pathogenesis is its ability to form complex biofilm structures. Pneumococci in a biofilm mode of growth display decreased sensitivity to antibiotics and increased resistance to host immune responses (6). These properties make the bacteria recalcitrant to treatment and highlight the need to better understand the molecular mechanisms that drive biofilm development. Activation of the competence pathway is critical for biofilm development. Previous *in vitro* studies have demonstrated that while cell-adherence and early biofilm formation are competence-independent, an intact competence system is required in the later stages of biofilm development. It was shown that the competence pathway positively influences structure and stability of late stage biofilms (12). However, the molecules downstream of competence activation by ComDE that regulate biofilm development remain poorly understood. In this study, we present BriC, a previously uncharacterized peptide, that we show is regulated by competence and plays a role in promoting biofilm development and nasopharyngeal colonization.

We have presented extensive evidence that *briC* is a competence regulated gene. We have shown that induction of *briC* is triggered by addition of CSP and requires ComE. Further, we have also shown that the *briC* promoter encodes the consensus ComE-binding box, and that *briC* expression follows the temporal pattern described for genes directly regulated by ComE. Previous studies have used microarray analysis to identify pneumococcal genes differentially regulated upon CSP stimulation (18,23) and have categorized these genes into two main categories - early genes regulated by ComE or late genes regulated by the alternative sigma factor, ComX. In their study, *briC* was found to be upregulated in a pattern consistent with early genes. However, the upregulation was not found to be statistically significant, and this study is the first validation of *briC* as a competence-regulated peptide.

We have provided evidence that *briC* stimulates biofilm development on abiotic surfaces and promotes nasopharyngeal colonization in a murine model. These findings are consistent with studies that show that pneumococcal biofilms contribute to colonization. Colonization of the upper respiratory tract is a requisite for pneumococcal dissemination to distant anatomical sites and subsequent disease (10). These sessile communities serve as a source of pneumococcal cells with an activated virulence-associated transcription program. That is, when compared to cells originating from a planktonic mode of growth, those originating from a biofilm mode of growth are more likely to cause disease upon infecting other tissues (11). In this manner, increased biofilm development likely heightens the risk for disease. Biofilms and competence are also associated with transformation efficiency. We have observed a mild but significant decrease in the transformation efficiency of *briC*-deletion mutants relative to WT R6D cells (**Fig. S1**). Finally, colonization of the upper respiratory tract is also a reservoir for pneumococcal transmission. Transmission occurs when cells migrate from the nasopharynx of one host to that of another. Thus, BriC’s contribution to colonization may influence both disease severity and transmission.

While it has been established that CSP contributes to biofilm development, the competence-dependent genes that regulate biofilm development are not well understood (7,12). Our finding that increased levels of *briC* can fully rescue biofilm defects from a *comE* deletion mutant *in vitro*, and partially rescue its colonization defects *in vivo* suggests that *briC* expression may bypass the requirement for competence in biofilm development. ComE is a key regulator of competence whose activity is required to regulate approximately 5-10% of the genome, and as such deletion of *comE* is expected to have substantial global consequences (18,23). In this context, it is noteworthy that overexpression of one gene (*briC*) in the *comE*-deletion mutant was able to improve colonization in the murine model. These findings strongly suggested that BriC is a molecular link between competence, biofilm development, and colonization.

Our data suggests that many strains have multiple inputs to the regulation of *briC*. Shared across all strains is the regulation by ComE, the key regulator of the competence pathway. Competence is responsive to environmental cues, such as changes in cell density, pH, mutational burden in cells, and exposure to antibiotics (16,51–53). Conversely, competence is inhibited by the degradation of CSP via the activity of the CiaHR TCS and the serine protease, HtrA (54,55). Factors altering competence will also alter *briC* levels due to its competence-dependent induction. Our comparative genomics suggest that a subset of pneumococcal lineages may encode an additional *briC*-regulatory element. Specifically, the *briC* promoter differs across strains, in that a subset of lineages encodes a long promoter with a RUP sequence (P*briC*_long_) and has higher basal levels of *briC* expression. This long promoter is constitutively active, even when competence is off.

The long promoter is encoded in the vast majority of strains from the PMEN1 lineage (Spanish-USA) and the PMEN14 (Taiwan-19F) lineages. These lineages are prominent in the clinical setting; they are multi-drug resistant and pandemic (56–58). This additional competence-independent regulation of the long promoter may provide promoter-binding sites for additional regulators or reflects consequences of positional differences for the existing promoter binding sites. Our biofilm and colonization experiments suggest that encoding the long *briC* promoter has functional consequences. We conclude that the response of *briC* to competence is ubiquitous, but that additional lineage-specific factors influence *briC* regulation and downstream phenotypic consequences.

We propose a model where *briC* encodes a signaling molecule with a role in biofilm development and colonization. First, the transcription of *briC* is induced by ComE through competence signal transduction pathway in all lineages, and possibly by additional regulator(s) in a subset of lineages. Once this Gly-Gly peptide is produced, we propose that it is exported through ABC transporters, a process in which ComAB plays a prominent role. Based on a bioinformatic comparison with other Gly-Gly peptides we suggest that BriC is cleaved into its active form (BRIC) during export. It is tempting to speculate that BRIC is a new member of the expanding set of pneumococcal secreted peptides that signal to neighboring cells promoting population-level behaviors. In this era of emerging antibiotic resistance, it is imperative that we test the potential of alternative strategies to inhibit bacterial carriage and disease. One such strategy is to specifically target bacterial communities and population-level behaviors. In that regard, molecules such as BriC present promising alternatives to be used as targets for discovery of novel drugs and therapeutic interventions.

## Materials & Methods

### Bacterial strains & growth conditions

Three wild-type (WT) *Streptococcus pneumoniae* strains were used for this experimental work. The majority of studies were performed on a penicillin-resistant derivative of R6 (R6D); this strain was generated from a cross where parental strain R6 was recombined with Hungary19A and the recombinant was selected for penicillin resistance (59). The *briC* allele in R6D is identical to the allele present in the parental R6. This laboratory strain is non-encapsulated and does not colonize mice, thus mice colonization experiments were performed with the serotype 2 D39 strains (GenBank CP000410)(60). The D39 strain contains the same *briC* allele as is present in the R6D strain, which has been used for most of the work in this study. Finally, for a representative of PMEN1, we used the carriage isolate, PN4595-T23 (GenBank ABXO01) graciously provided by Drs. Alexander Tomasz and Herminia deLancastre (61).

Colonies were grown from frozen stocks by streaking on TSA-II agar plates supplemented with 5% sheep blood (BD BBL, New Jersey, USA). Colonies were then used to inoculate fresh Columbia broth (Remel Microbiology Products, Thermo Fisher Scientific, USA) and incubated at 37°C and 5% CO_2_ without shaking. When noted, colonies were inoculated into acidic Columbia broth prepared by adjusting the pH of Columbia broth to 6.6 using 1M HCl. Acidic pH was used to inhibit the endogenous activation of competence.

### Construction of mutants

The mutant strains (R6DΔ*briC and* PN4595Δ*briC*) were constructed by using site-directed homologous recombination to replace the region of interest with erythromycin-resistance gene (*ermB*) or kanamycin-resistance gene (*kan*), respectively (**Table S4**). *Kan* and spectinomycin-resistance gene (*aad9*) were used to construct ∆*comE* strains in R6D and PN4595-T23 respectively. Briefly, the transformation construct was generated by assembling the amplified flanking regions and antibiotic resistance cassettes. ~2kb of flanking regions upstream and downstream of the gene of interest was amplified from parental strains by PCR using Q5 2x Master Mix (New England Biolabs, USA). The antibiotic resistance genes, *kan* and *aad9* were amplified from *kan*-*rpsL* Janus Cassette and pR412, respectively (provided by Dr. Donald A. Morrison), and *ermB* was amplified from *S. pneumoniae* SV35-T23. SV35-T23 is resistant to erythromycin because of the insertion of a mobile element containing *ermB* (61). These PCR fragments were then assembled together by sticky-end ligation of restriction enzyme-cut PCR products. The deletion mutant in R6D is an overexpressor of the downstream peptide (*spr_0389*).

The *briC* complemented and overexpressor strains were generated using constructs containing the CDS of *briC* along with either its entire native promoter region or overexpressing promoter respectively, ligated at its 3’ end with a kanamycin resistance cassette. The promoters used to overexpress *briC* included either the constitutive *amiA* promoter, or P*briC*_long_. These were assembled with the amplified flanking regions by Gibson Assembly using NEBuilder HiFi DNA Assembly Cloning Kit. The construct was introduced in the genome of R6D downstream of the *bga* region (without modifying *bga*), a commonly employed site (62). Primers used to generate the constructs are listed in **Table S5**. Like R6DΔ*briC*, R6DΔ*briC::briC* is also an overexpressor of the downstream peptide (*spr_0389*), which is annotated as a pseudogene in strains R6D, R6 and D39 (**Fig. 2A**). The expression of *spr_0389* remains unchanged in the mutant and the complement (data not shown from qRT-PCR).

The R6DΔ*comE::*P*briC*_long_-*briC* strain was constructed by replacing *comE* with spectinomycin resistant cassette in the R6D P*briC*_long_-*briC* strain. *comAB*-deletion mutant in a *briC* overexpressor R6D genomic background strain (R6DΔ*comAB::*P*briC*_long_-*briC*) was constructed by transforming the R6D::P*briC*_long_-*briC* strain with the genomic DNA of ADP226. ADP226 is a strain from the D39 genomic background with *comAB* replaced by erythromycin resistance cassette. To make the construct, the flanking regions and erythromycin resistance cassette were amplified, and then assembled together by sticky-end ligation of restriction enzyme-cut PCR products. The construct was then transformed into D39 ADP225 (unpublished) and selected on Columbia blood agar supplemented with 0.25 μg mL^−1^ erythromycin.

The *briC* promoter region was modified by shuffling the ComE-binding box (R6DΔ*briC::*P*briC*_Shuffled ComE-box_*-briC*). The ComE-binding box was shuffled using PCR by amplifying from R6DΔ*briC::briC* and introducing the shuffled sequence (CAGACCAGTTAGTCTAGGATAGAGCTTAAG) into the primers. The resulting amplicons were assembled using Gibson Assembly. The modified construct was transformed into R6DΔ*briC* strain in the region downstream of the *bgaA* gene.

The D39 *briC* deletion mutant (D39Δ*briC*), *briC* complemented (D39Δ*briC::briC*), *comE* deletion mutant (D39Δ*comE*), *briC* overexpressor in *comE* deletion background (D39Δ*comE*::P*briC*_long_-*briC*), and *briC* expressed from a promoter with a shuffled ComE-binding box (R6DΔ*briC::*P*briC*_Shuffled ComE-_box-*briC*) strains were generated by transformation with the corresponding constructs amplified from R6D into strain D39.

### Construction of *lacZ* fusions

Chromosomal transcriptional *lacZ*-fusions to the target promoters were generated to assay promoter activity. These *lacZ*-fusions were generated via double crossover homologous recombination event in the *bgaA* gene using modified integration plasmid pPP2. pPP2 was modified by introducing *kan* in the multiple cloning site, in a direction opposite to *lacZ*. The modified pPP2 was transformed into *E. coli* TOP10. The putative *briC* promoter regions were amplified from R6 and PN4595-T23 strains, and modified to contain KpnI and XbaI restriction sites, which were then assembled in the modified pPP2 plasmid by sticky-end ligation of the enzyme digested products. These plasmids were transformed into *E. coli* TOP10 strain, and selected on LB (Miller’s modification, Alfa Aesar, USA) plates, supplemented with ampicillin (100µg/ml). These plasmids were then purified by using E.Z.N.A. Plasmid DNA Mini Kit II (OMEGA bio-tek, USA), and transformed into pneumococcal strains R6 and PN4595-T23 and selected on Columbia agar plates supplemented with kanamycin (150µg/ml).

### Bacterial transformations

For all bacterial transformations to generate mutants, target strains (R6D or D39) were grown in acidic Columbia broth, and 1µg of transforming DNA along with 125µg/mL of CSP1 (sequence: EMRLSKFFRDFILQRKK; purchased from GenScript, NJ, USA) was added to them when the cultures reached an OD_600_ of 0.05, followed by incubation at 37°C. After 2 hours, the treated cultures were plated on Columbia agar plates containing the appropriate antibiotic; erythromycin (2µg/ml), or kanamycin (150µg/ml). Resistant colonies were cultured in selective media, and the colonies confirmed using PCR. Bacterial strains generated in this study are listed in **Table S4**.

For transformation efficiency experiments, R6D strain was grown in acidic Columbia broth until it reached an OD_600_ of 0.05. At this point, number of viable cells was counted by plating serial dilutions on TSA-blood agar plates. Transformations were carried out by adding either 100ng or 500ng of transforming DNA in the media supplemented with 125µg/mL of CSP1 and incubated at 37°C for 30mins. For transforming DNA, we used either genomic DNA or PCR products. The donor DNA contained spectinomycin-resistance gene (*aad9*) in the inert genomic region between *spr_0515* and *spr_0516*. This construct was generated in PN4595-T23, spec^R^, followed by its amplification and transformation into R6D and Taiwan-19F strains (Sp3063-00). The genomic DNA was extracted from Taiwan-19F, spec^R^ strain. The purified linear DNA was an amplimer of the region from R6D. After 30 minutes, the cultures were plated on Columbia agar plates containing spectinomycin (100µg/ml), incubated overnight, and colonies were counted the next day.

### RNA extraction

RNA extraction consists of sample collection, pneumococcal cell lysis, and purification of RNA. For qRT-PCR analysis, the strains (R6D and R6DΔ*comE*) were grown to an OD_600_ of 0.3 in acidic Columbia broth, followed by CSP1 treatment for 0, 10, or 15 minutes. For *in vitro* transcriptomic analysis using NanoString Technology, the R6D strain was grown to an OD_600_ of 0.1 in Columbia broth (in one experimental set, the samples were grown in sub-lethal concentration of penicillin (0.8µg/ml) for an hour). RNA was collected in RNALater (Thermo Fisher Scientific, USA) to preserve RNA quality and pelleted. For the *in vivo* experiments, the RNA was extracted from middle-ear chinchilla effusions infected with PN4595-T23 and PN4595-T23Δ*comE* strains, and preserved by flash freezing the effusion. In all the samples, the pneumococcal cell lysis was performed by re-suspending the cell pellet in an enzyme cocktail (2mg/ml proteinase K, 10mg/ml lysozyme, and 20µg/ml mutanolysin), followed by bead beating with glass beads (0.5mm Zirconia/Silica) in FastPrep-24 Instrument (MP Biomedicals, USA). Finally, RNA was isolated using the RNeasy kit (Genesee Scientific, USA) following manufacturer’s instructions. For analysis with the NanoString, which does not require pure DNA, the output from the RNeasy kit was loaded on the machine without further processing. For analysis using qRT-PCR, contaminant DNA was removed by treating with DNase (2U/µL) at 37°C for at least 45 mins. The RNA concentration was measured by NanoDrop 2000c spectrophotometer (Thermo Fisher Scientific, USA) and its integrity was confirmed on gel electrophoresis. The purity of the RNA samples was confirmed by the absence of a DNA band on an agarose gel obtained upon running the PCR products for the samples amplified for *gapdh*.

### NanoString Technology for transcriptional analysis

nCounter Analysis System from NanoString Technology provides a highly sensitive platform to measure gene expression both *in vitro* and *in vivo*, as previously described (63). Probes used in this study were custom-designed by NanoString Technology, and included housekeeping genes *gyrB* and *metG*, which were used as normalization controls. 5µL of extracted RNA samples were hybridized onto the nCounter chip following manufacturer’s instructions. RNA concentration ranged from 80-200ng/µL for *in vivo* samples, and between 60-70ng total RNA for *in vitro* samples. A freely available software from manufacturers, nSolver, was used for quality assessment of the data, and normalization. The RNA counts were normalized against the geometric mean of *gyrB* and *metG* (64,65). The 16S rRNA gene is not optimal for normalization in the NanoString, as the high abundance of this transcript packs the field of view. Pearson’s Correlation Coefficient was used to estimate correlation in the expression levels of different genes.

### qRT-PCR for transcriptional analysis

High quality RNA was used as a template for first-strand cDNA synthesis SuperScript VILO synthesis kit (Invitrogen). The resulting product was then directly used for qRT-PCR using PerfeCTa SYBR Green SuperMix (Quantabio, USA) in an Applied Biosystems 7300 Instrument (Applied Biosystems, USA).16S rRNA counts were used for normalization. The raw data was then run through LinregPCR for expression data analysis, where the output expression data is displayed in arbitrary fluorescence units (N_0_) that represent the starting RNA amount for the test gene in that sample (66,67). Fold-change relative to WT was then calculated for each individual experiment.

### β-galactosidase assay

β-galactosidase assays were performed as previously described (68) using cells that were grown in acidic Columbia broth to exponential phase. Cells were either left untreated, or independently treated with CSP1 (EMRLSKFFRDFILQRKK) or CSP2 (EMRISRIILDFLFLRKK) (Genscript, USA) for 30 minutes and processed for analysis.

### Biofilm formation assay

Pneumococcal cultures grown in Columbia broth were used to seed biofilms on abiotic surfaces. When the cultures reached an OD_600_ of 0.05, each bacterial strain was seeded on 35MM glass bottom culture dishes (MatTek Corporation, USA). To promote biofilm growth, the plates were incubated at 37°C and 5% CO_2_. Every 24 hours, the supernatant was carefully aspirated, followed by addition of the same volume of pre-warmed Columbia broth at one-fifth concentration. The biofilm samples were fixed at two time-points: 24 and 72 hrs. For fixing, the supernatants were carefully aspirated, and biofilms were washed thrice with PBS to remove non-adherent and/or weakly adherent bacteria. Subsequently, biofilms were fixed with 4% PFA (Electron Microscopy Sciences), washed three times with PBS, and prepared for confocal microscopy.

### Confocal microscopy & quantification of biofilms

Fixed biofilms were stained with SYTO59 Nucleic Acid Stain (Life Technologies, USA) for 30 minutes, washed three times, and preserved in PBS buffer for imaging. Confocal microscopy was performed on the stage of Carl Zeiss LSM-880 META FCS, using 561nm laser line for SYTO59 dye. Stack were captured every 0.46 µm, imaged from the bottom to the top of the stack until cells were visible, and reconstructed in Carl Zeiss black edition and ImageJ. The different biofilm parameters (biomass, maximum thickness, and average thickness over biomass) were quantified using COMSTAT2 plug-in available for ImageJ (69). For depiction of representative reconstructed Z-stacks, empty slices were added to the images so the total number of slices across all the samples were the same. These reconstructed stacks were pseudo-colored according to depth using Carl Zeiss black edition. The color levels of the images being used for representation purposes were adjusted using GNU Image Manipulation Program (GIMP).

### Testing peptide secretion using the Nano-Glo HiBiT extracellular detection system

HiBiT constructs were designed by fusing the C-terminus of the region of interest with the 11-amino acid HiBiT peptide using a 10-amino acid linker. The region of interest was the putative secretion signal (until the double glycine) of the *briC* gene. The expression of these constructs was designed to be controlled by the *briC* promoter region. The construct was introduced in the genome of R6D and R6DΔ*comAB* strains downstream of the *bgaA* gene (without modifying *bgaA*).

R6D strains containing HiBiT constructs were started from overnight blood agar plates into acidic Columbia broth (pH 6.6) and incubated at 37°C and 5% CO_2_ without shaking. Cultures were grown to an OD_600_ of ~0.2. Cultures were either left untreated or treated with 125µg/mL of CSP1 for 30 minutes, followed by measuring optical density at 600nm. Cells were pelleted by centrifuging the cultures for 5 minutes at 3700_×g_. The resulting supernatants were removed and filtered through 0.2μm syringe filters. The cell pellets were resuspended in equal volume of PBS. To obtain cell lysate, triton X-100 was added to 1ml of the resuspended cells to a final concentration of 1%. Additionally, to minimize non-specific binding, triton X-100 was also added to 1ml of the filtered supernatant to a final concentration of 1%. 75μl of the supernatant, whole cells, lysates, were added to a Costar96 well flat white tissue culture treated plates and mixed with an equal volume of the Nano-Glo Extracellular Detection System reagent as specified in the manufacturer’s instructions. Additionally, media and PBS samples were used as controls. Reactions were incubated at room temperature for 10 minutes followed by measuring luminescence on a Tecan Spark with an integration time of 2000 milliseconds.

### *In vivo* transcriptomic analysis using chinchilla OM model

All chinchilla experiments were conducted with the approval of Allegheny-Singer Research Institute (ASRI) Institutional Animal Care and Use Committee (IACUC) A3693-01/1000. Research grade young adult chinchillas (*Chinchilla lanigera*) weighing 400-600g were acquired from R and R Chinchilla Inc., Ohio. Chinchillas were maintained in BSL2 facilities and experiments were done under subcutaneously injected ketamine-xylazine anesthesia (1.7mg/kg animal weight for each). Chinchillas were infected with 100 CFUs in 100µL of *S. pneumoniae* PN4595-T23 by transbullar inoculation within each middle ear. For RNA extraction, chinchillas were euthanized 48h post-inoculation of pneumococcus, and a small opening was generated through the bulla to access the middle ear cavity. Effusions were siphoned out from the middle ear and flash frozen in liquid nitrogen to preserve the bacterial RNA. Animals were euthanized by administering an intra-cardiac injection of 1mL potassium chloride after regular sedation.

### Murine colonization model

The role of *briC* in experimental pneumococcal colonization was assessed as previously described (70,71). For this, 10 weeks old female CD1 mice (Charles River), weighing approximately 30-35 g were anesthetized with 2.5% isoflurane over oxygen (1.5 to 2 liter/min), and administered intranasally with approximately 1X10^5^ CFU/mouse in 20µl PBS. At predetermined time intervals, a group of 5 mice were euthanized by cervical dislocation, and the nasopharyngeal lavage of each animal was obtained using 500µl PBS. The pneumococci in nasopharyngeal wash were enumerated by plating the serial dilutions onto blood agar plates.

### Statistical tests

The statistical differences among different groups were calculated by performing ANOVA followed by Tukey’s post-test, unless stated otherwise. p-values of less than 0.05 were considered to be statistically significant.

### Distribution of *briC* across streptococcal strains

To identify *briC* homologs we used tBLASTn with default parameters on the RAST database to search the genome sequences of all fifty-five strains. Predicted protein sequences were downloaded as well as nucleotide sequences for the *briC* homolog and 1500-bp flanking regions surrounding the *briC* homolog. Predicted protein sequences for BriC were aligned using NCBI Cobalt (72) and visualized using Jalview (73). One sequences (CDC3059-06) appeared to have a frame-shift after a string of guanines. Given that sequencing technologies are often inaccurate after a string of identical bases, we curated this sequence in the dataset. The sequences were translated in Jalview, and organized based on polymorphisms in the translated sequences.

The *briC* alleles were then organized in the context of the species tree. For this we used a published phylogenetic tree (34,38). As previously described, the whole genome sequence (WGS) for these strains were aligned using MAUVE (74,75), the core region was extracted and aligned using MAFFT (FFT-NS-2) (76). Model selection was performed using MODELTEST (77), and the phylogenetic tree was built with PhyML 3.0 (78), model GTR+I(0.63) using maximum likelihood and 100 bootstrap replicates. On the visualization, each allelic type is shape-coded, and the visualization was generated using the Interactive Tree of Life (iTOL) (79).

Next, we expanded the search to a set of 4,034 genomes. These correspond to the genomes within pubMLST, with at least 2Mb of genomic data (Genome IDs are listed in **Table S6**). We used BLASTn to search for genomes that encode sequences that are at least 70% identical over 70% of their length to *briC* alleles 1A or 1B. The 3,976 hits were organized to parse out and enumerate the unique sequences using Python. Next, the hits were visualized and further annotated using Jalview (73). As in the smaller genomic set, one allele representative appeared to have a frame-shift after a string of guanines and was curated in the dataset. Next the DNA sequences were translated, and the predicted protein sequences were organized to display the unique alleles. The resulting 19 coding sequence were colored in Jalview based on percent identity to highlight the variability (**Fig. 2B**). To search for *briC* in related species, we performed a BLASTp analysis in NCBI. We used alleles 1A and 1B as query sequences, default parameters, and the non-redundant database excluding *Streptococcus pneumoniae* (taxid: 1313).

### Analysis of *briC* promoter region

In order to examine the structure of the promoter region upstream of the *briC* gene, a 1500-bp flanking region on both sides of the *briC* gene was pulled from the RAST database (80). Sequences were aligned using Kalign (81) and then visualized with Jalview (73). The alignment revealed two clear groups within the dataset: those with the RUP insertion and those without. We also noted that CDC1087-00 may have an additional mobile element inserted within the RUP. However, given that the RUP and this mobile element exist in multiple places in the genome, we cannot determine whether this is real or an artifact of assembly without the isolate. Thus, we opted not to use the promoter sequence for the consensus in Figure 2A, and we did not mark this genome as having a long promoter in Figure 4A. We marked the species tree with allelic variants that contain the RUP insertion. We observed that RUP was present in the representative isolates from two clinically important lineages PMEN1 and PMEN14. To check the distribution of the long promoter in a larger set strains, we used PubMLST (39) to inspect 4,034 sequences with complete genomes. The sequence IDs for these 4,034 sequences are listed in **Table S6**. This set includes 198 ST81 (PMEN1), as well as 104 ST236 (PMEN14) and 15 ST320 (PMEN14) strains. For analysis of the ComE-binding box, the ComE consensus sequence was extracted from the promoter regions of the pneumococcal strains and aligned with Jalview. The logo was generated using WebLogo (82).

### Ethics statement

Mouse experiments were performed at the University of Leicester under appropriate project (permit no. P7B01C07A) and personal licenses according to the United Kingdom Home Office guidelines under the Animals Scientific Procedures Act 1986, and the University of Leicester ethics committee approval. The protocol was agreed by both the U.K. Home Office and the University of Leicester ethics committee. Where specified, the procedures were carried out under anesthetic with isoflurane. Animals were housed in individually ventilated cages in a controlled environment, and were frequently monitored after infection to minimize suffering. Chinchilla experiments were performed at the Allegheny-Singer Research Institute (ASRI) under the Institutional Animal Care and Use Committee (IACUC) permit A3693-01/1000. Chinchillas were maintained in BSL2 facilities, and all experiments with chinchillas were done under subcutaneously injected ketamine-xylazine anesthesia (1.7mg/kg animal weight for each). All chinchillas were maintained in accordance with the applicable portions of the Animal Welfare Act, and the guidelines published in the DHHS publication, Guide for the Care and Use of Laboratory Animals.

## Acknowledgements

We thank Drs. Alexander Tomasz and Herminia deLancastre for the PMEN1 strain PN4595-T23 and the R6D strain used in this study. We would also like to thank Dr. Donald A. Morrison for *kan-rpsL* Janus cassette and the plasmid pR412. We thank Anagha Kadam for help on data analysis, Rolando A. Cuevas for help analyzing biofilm images, and Emilio I. Rodriguez for his support with experiments. We truly appreciate the insight provided by Dr. Suzanne Dawid and Charles Wang regarding the HiBiT assay to test the role of ComAB in BriC export.

## Supplementary Result

### Examining the role of *briC* in transformation

One of the main phenotypic consequences of the competence pathway is transformation. Since ComE regulates the expression of *briC*, we investigated whether *briC* plays a role in regulating transformation efficiency. To this end, we added different amounts (100ng or 500ng) of exogenous DNA (genomic or amplified linear fragments) to pneumococcal cells. We found a minor decrease in the transformation efficiency of the *briC-*deletion mutant cells (R6DΔ*briC*) relative to the WT cells (**Fig. S1**). A strain with complemented *briC* (R6DΔ*briC*::*briC*) exhibited a partial restoration of this defect. A strain with overexpression of *briC* (R6DΔ*briC*::*briC*-OE) displayed a greater restoration of the transformation efficiency, as compared to R6DΔ*briC*::*briC* cells, but not a full rescue (**Fig. S1B**). These results suggest that *briC* may play a role in regulating transformation efficiency.

## Supplementary Tables

Table S1: Strains used in genomic comparisons and phylogenetic tree.

Table S2: CSP pherotypes and *briC* alleles for pneumococcal genomes in figure 4A.

Table S3: HiBiT assay measuring bioluminiscence of the construct containing BriC leader fused with HiBiT, both in the absence and presence of CSP in supernatants, whole cell surfaces and lysates.

Table S4: Strains used in this experimental work.

Table S5: Primers used in this study.

Table S6: List of the 4,034 isolates used in this study. These numbers correspond to isolate IDs of strains in pubMLST.

## Supplementary Files

File S1: Representative *briC* alleles from *Streptococcus* sp.

File S2: Representative sequences of the short and long *briC* promoter regions with the RUP highlighted in bold.

